# UNSEG: unsupervised segmentation of cells and their nuclei in complex tissue samples

**DOI:** 10.1101/2023.11.13.566842

**Authors:** Bogdan Kochetov, Phoenix Bell, Paulo S. Garcia, Akram S. Shalaby, Rebecca Raphael, Benjamin Raymond, Brian J. Leibowitz, Karen Schoedel, Rhonda M. Brand, Randall E. Brand, Jian Yu, Lin Zhang, Brenda Diergaarde, Robert E. Schoen, Aatur Singhi, Shikhar Uttam

## Abstract

Multiplexed imaging technologies have made it possible to interrogate complex tumor microenvironments at sub-cellular resolution within their native spatial context. However, proper quantification of this complexity requires the ability to easily and accurately segment cells into their sub-cellular compartments. Within the supervised learning paradigm, deep learning based segmentation methods demonstrating human level performance have emerged. However, limited work has been done in developing such generalist methods within the label-free unsupervised context. Here we present an unsupervised segmentation (UNSEG) method that achieves deep learning level performance without requiring any training data. UNSEG leverages a Bayesian-like framework and the specificity of nucleus and cell membrane markers to construct an *a posteriori* probability estimate of each pixel belonging to the nucleus, cell membrane, or background. It uses this estimate to segment each cell into its nuclear and cell-membrane compartments. We show that UNSEG is more internally consistent and better at generalizing to the complexity of tissue morphology than current deep learning methods. This allows UNSEG to unambiguously identify the cytoplasmic compartment of a cell, which we employ to demonstrate its use in an exemplar biological scenario. Within the UNSEG framework, we also introduce a new perturbed watershed algorithm capable of stably and automatically segmenting a cluster of cell nuclei into individual cell nuclei that increases the accuracy of classical watershed. Perturbed watershed can also be used as a standalone algorithm that researchers can incorporate within their supervised or unsupervised learning approaches to extend classical watershed, particularly in the multiplexed imaging context. Finally, as part of developing UNSEG, we have generated a high-quality annotated gastrointestinal tissue (GIT) dataset, which we anticipate will be useful for the broader research community. We demonstrate the efficacy of UNSEG on the GIT dataset, publicly available datasets, and on a range of practical scenarios. In these contexts, we also discuss the possibility of bias inherent in quantification of segmentation accuracy based on *F*_1_ score. Segmentation, despite its long antecedents, remains a challenging problem, particularly in the context of tissue samples. UNSEG, an easy-to-use algorithm, provides an unsupervised approach to overcome this bottleneck, and as we discuss, can help improve deep learning based segmentation methods by providing a bridge between unsupervised and supervised learning paradigms.

## Introduction

Recent innovations in highly multiplexed immunofluorescence imaging^1–15^ have substantially increased the range of antigens that can be spatially profiled in a tissue sample, from 3 − 5 targets to ∼ 60^16^. Segmentation is a required step for quantitatively associating their spatial expressions with individual cells. Since 2012, when AlexNet^17^, a deep convolutional neural network (CNN), outperformed other methods in the ImageNet classification challenge, there has been a paradigm shift towards using CNN based deep learning (DL) frameworks^18^ trained on curated datasets for cell and nucleus segmentation tasks^19–28^. Among them, Cellpose^25^ − a DL method based on a U-Net architecture utilizing gradient flow representation of cells− and Mesmer^26^ − a DL method based on ResNet50 architecture − have demonstrated human-level performance in the highly multiplexed imaging context. However, due to their dependence on stochastic gradient descent and back-propagation based optimization during the training step, it remains difficult to identify the contribution of each neuron to the eventual segmentation outcome, and as a consequence explain the source of errors in segmentation when they occur^29^. As a result, improving performance of these black-box DL models requires rewiring the input-output mapping via training on additional datasets^30^. However, in complex tissue samples with considerable heterogeneity and ambiguity in cellular organization, it is unclear whether retraining alone will consistently improve results across all samples, or if multiple DL models need to be constructed and used through a trial and error approach, with the hope that their performance will optimally generalize. Curation of accurately annotated datasets of sufficient quality that capture the tissue microenvironment diversity also remains a critical challenge.

In contrast to DL approaches, most unsupervised cell segmentation methods^31–43^ do not require training data, are explainable, and therefore where needed, can be optimized for individual images. However, to the best of our knowledge, to date no unsupervised segmentation method capable of approaching DL method performance exists, or has even been considered feasible. Here, we present a new unsupervised segmentation algorithm (UNSEG) capable of performing sub-cellular segmentation of tissue sample images with accuracy on par with state-of-the-art DL segmentation approaches such as Cellpose and Mesmer. UNSEG achieves this performance in two stages. At the first stage, UNSEG quantifies the intrinsic contrast provided by any nucleus and cell membrane specific markers at the local and global scale, and jointly exploits it to assign each pixel to the nucleus, cell membrane, or the background class. This pixel assignment is implemented with the help of Bayesian-like framework that computes *a priori* distributions and an image contrast-based likelihood function to estimate the posterior probabilities of each pixel belonging to nucleus, cell membrane or background classes. UNSEG uses the posterior probabilities to assign the pixel to the correct compartment. At the second stage, it parses the semantic pixel assignments into topologically consistent nuclei and cells. Towards this goal UNSEG introduces perturbed watershed, a new algorithm we have developed, to correctly partition a nucleus cluster into individual nuclei. The final output of UNSEG are nucleus and cell segmentations corresponding to the input image.

We have curated a labeled gastrointestinal tissue (GIT) dataset comprising of diverse images of gastrointestinal tissue to benchmark UNSEG performance. We anticipate that this dataset will also be useful to DL researchers and the broader research community and help ameliorate shortage in annotated imaging datasets^30^. We have also tested UNSEG performance on public datasets, with images drawn from diverse tissue types and diseases beyond the gastrointestinal system, that have been labeled with different nucleus and cell membrane markers and acquired at different magnifications and resolutions. Additionally, we also demonstrate applicability of UNSEG in a variety of real-world cases that include, weakly expressing markers, non-specific markers, different nucleus markers, and multiplexed ion beam imaging (MIBI). In the context of these diverse scenarios, we also discuss how quantification of segmentation accuracy can potentially be biased depending on the nature of deviation of segmentation mask from the ground truth. Finally, we note that since UNSEG does not require any training data to segment tissue images, it can be used to generate high quality segmentation of unlabeled tissue images, which is majority of the data in real-world settings, as optimized initial estimates for improving DL models within unsupervised and semi-supervised settings. UNSEG, therefore, is an easy to use method for unsupervised sub-cellular segmentation of images of complex tissue samples that does not require extensive setup and performs on par with state-of-the-art DL methods. It also has the potential to improve the state-of-the-art in deep learning.

## Results

### UNSEG principle and design

Segmenting cells and nuclei in 2D images of tissue samples is challenging because of their complex morphology, ambiguous overlaps, and heterogeneity in spatial distribution of nucleus and cell-membrane markers within each cell. In the morphological context, although cells and their nuclei exhibit an overall convex topology, they locally deviate from it to varying degrees depending on cell types, and particularly in tumors with irregularly shaped cancer cells. Additionally, many cells in a tissue-dependent manner are clumped in clusters where their shape and overlap is difficult to parse. Cells in tissues also exhibit uneven intra-cellular distribution of marker expression. Together, these degrees of complexity make it difficult to consistently segment cells and nuclei using unsupervised segmentation approaches such as classical watershed^31,32,38^, shape and intensity prior^36,37,39–41^, and tracking of diffused gradient flow^33,34^, which have primarily been developed for segmenting cells in culture that lack tissue associated heterogeneity related to cellular morphology, expression and overlap. UNSEG framework overcomes these limitations by jointly exploiting the expression-based topology and distribution of markers specific to nuclei and cell-membranes (Figure 1). Such markers are also used in the supervised context of DL methods such as Cellpose and Mesmer.

**Figure 1.**
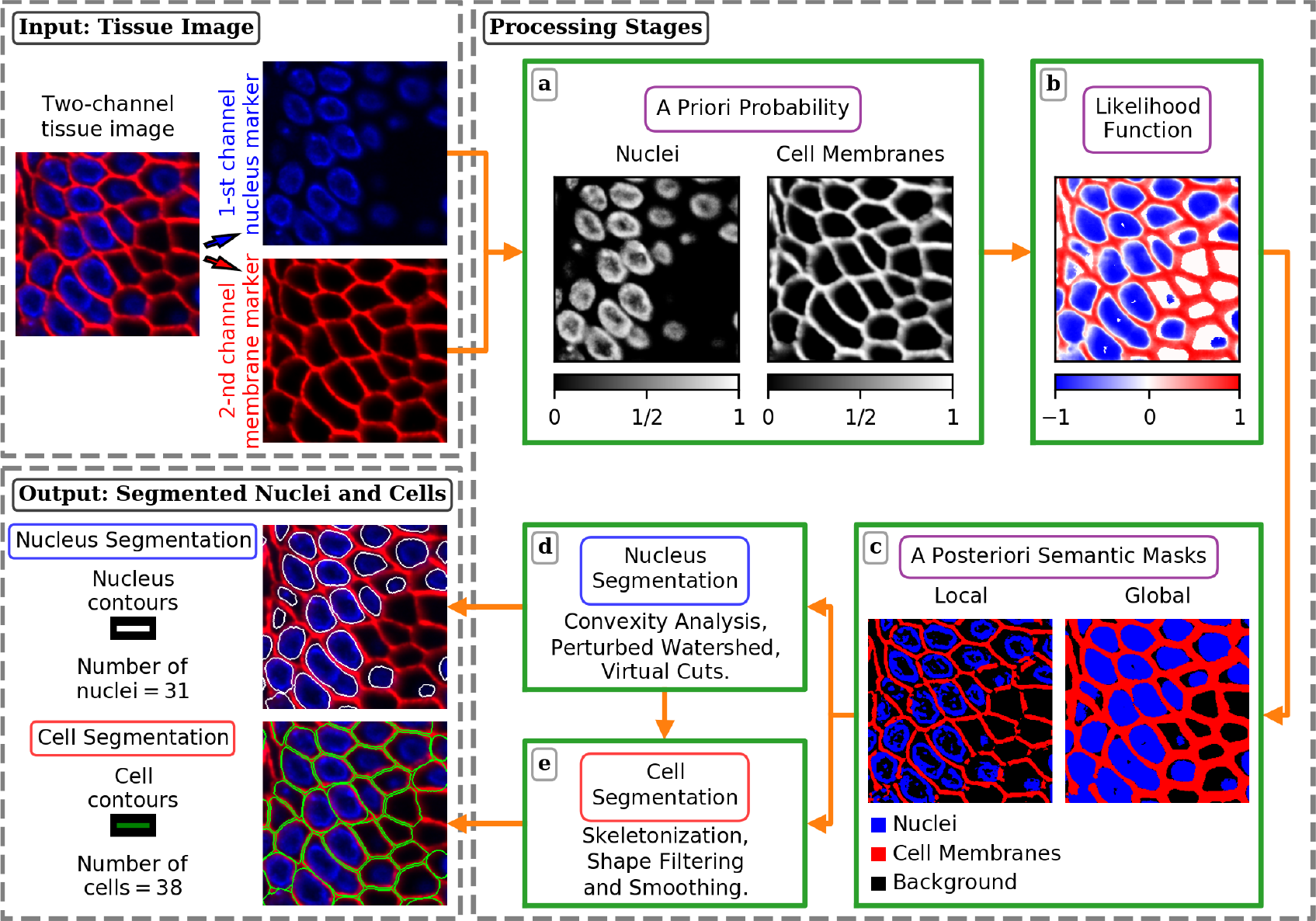
UNSEG framework. Input is a two-channel image comprising of nucleus (channel 1) and cell membrane (channel 2) marker expressions. **a** *A priori* spatial probability distributions of nucleus and cell membrane marker expressions. **b** Likelihood map of a pixel to belong to the nucleus or cell-membrane, quantified through the visual contrast function, which mimics human perception. **c** *A posteriori* local and global semantic segmentation masks respectively capturing local morphological heterogeneity and global nucleus and cell-membrane topology. **d** Instance segmentation of nuclei from semantic segmentation masks. **e** Instance segmentation of cells based on individual segmented nuclei and semantic masks. Nucleus and cell segmentation results of **d** and **e** form the UNSEG output. See Methods for more details.

UNSEG combines *a priori* probability of each image pixel belonging to a nucleus or cell-membrane (Figure 1a) with a contrast-based likelihood function (Figure 1b), to compute *a posteriori* semantic segmentation of image pixels into nucleus and cell-membrane (Figure 1c). UNSEG performs this segmentation both at the global level of the entire image, and at the local level in a neighborhood around each pixel (Figure 1c). The local segmentation captures the local heterogeneity in nucleus and cellular morphology, while the global segmentation ensures that the overall topological structure of the nuclei and cell membranes is preserved across the entire image. The final step of UNSEG utilizes these local and global nucleus and cell semantic masks to obtain instance segmentation of individual nuclei (Figure 1d) and cells (Figure 1e). This step includes partitioning nucleus clusters into individual nuclei based on convexity analysis, perturbed watershed and its ancillary function we refer to as virtual cuts. The latter two are briefly described below. The details of each step are described in Methods section. *Perturbed watershed*. Classical watershed based segmentation^44,45^ identifies individual nuclei in a cluster as watersheds, with each watershed basin representing a nucleus in the cluster. However, heterogeneity in spatial distribution of nucleus marker can make it difficult to uniquely identify the individual basins. Cellpose overcomes this problem in the supervised context by developing a gradient flow field representation of each nucleus whose ground truth is annotated by a human user^25^. This representation provided a stable and unique representation of nucleus basins. In the unsupervised context, we have developed a perturbed watershed approach (Figure 2 and Methods), where the initial watershed based segmentation (Figure 2i) of the nucleus cluster into individual nuclei is perturbed (Figures 2j-2m) based on an adaptive distance-transform estimate (Figure 2h) computed from the global nucleus cluster (Figure 2d) and local topology of the cell-membrane network (Figure 2e). Nuclei that are correctly segmented remain stable to the perturbations, while spuriously segmented nuclei collapse to a point-like object with area not exceeding a few pixels. When applied recursively, perturbed watershed partitions the nucleus cluster into individual nuclei. An example of a two-nuclei cluster is shown in Figure 2. Initial watershed partitions the cluster into three nuclei (Figure 2i), one of which shrinks to a point object on perturbation of the watershed seed point. The perturbation is performed in four directions: up, down, left, and right. In this example, the unstable nucleus collapsed for three of those perturbations (up, down, and left), indicating that the seed point is unstable and the corresponding segmentation is a spurious nucleus. Therefore, it is removed and the correct watershed based segmentation (Figure 2n) is obtained using the two remaining stable seed points and the original distance transform (Figure 2g). We note that perturbed watershed algorithm does not make any assumptions specific to the fluorescence based imaging modality. It is, in fact, agnostic to the imaging modality being used, and can be used to improve classical watershed results, wherever, the latter method is applicable.

**Figure 2.**
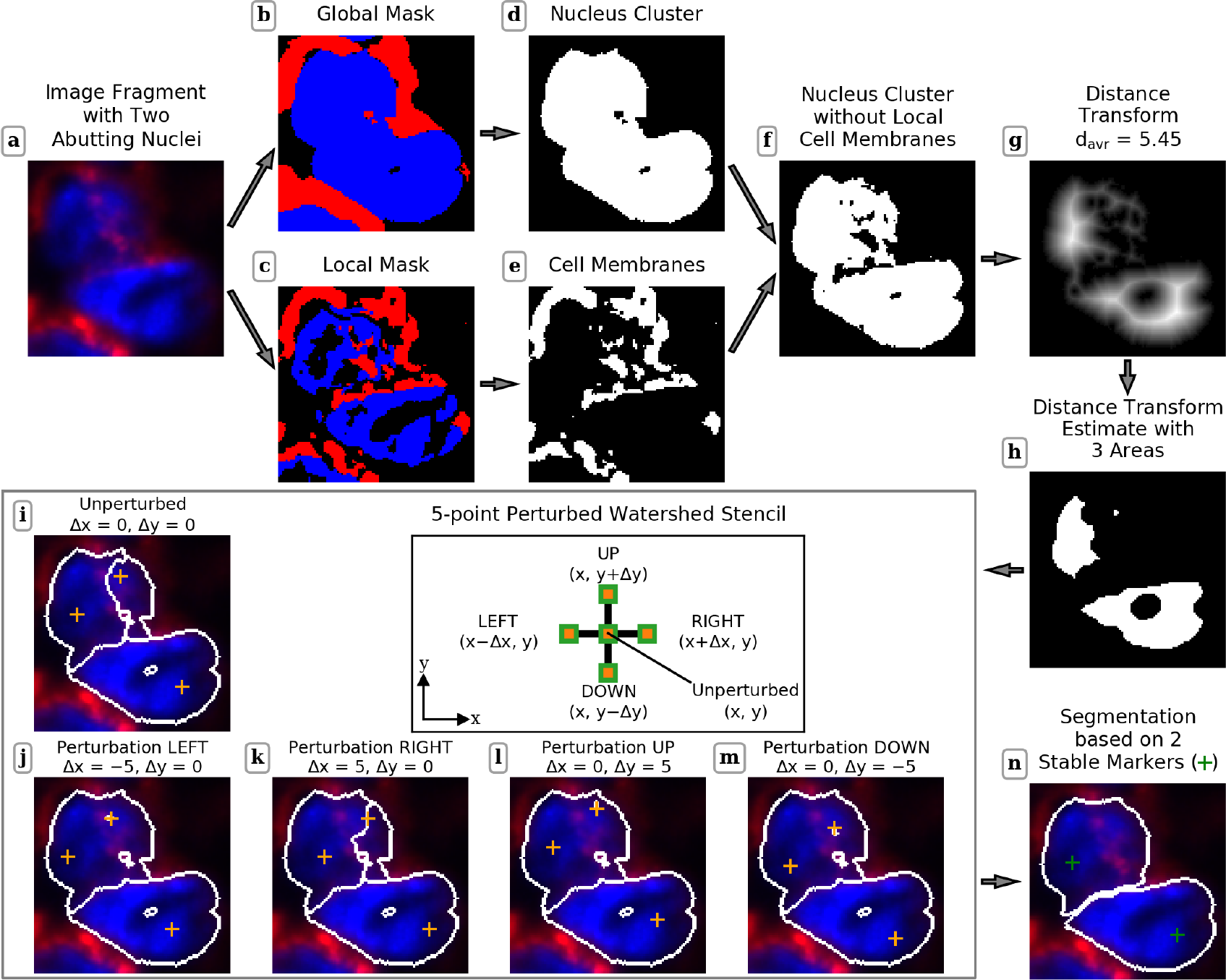
Perturbed watershed method. **a** Input image fragment with two abutting nuclei. **b-e** The posterior global and local masks of the input image from which the global nucleus cluster mask and local cell-membrane mask are extracted for downstream perturbed watershed analysis. **f** Global nucleus cluster mask with cuts corresponding to the local cell-membrane mask. **g** Distance transform of this modified global nucleus cluster mask. **h** Adaptive distance-transform estimate obtained by thresholding the distance transform by *d*_*avr*_ (see Methods). **i** Initial (unperturbed) watershed segmentation. **j-m** Perturbed watershed segmentations computed after shifting all markers from their unperturbed positions to the left (Δ*x* = −⌊*d*_*avr*_⌋), right (Δ*x* = ⌊*d*_*avr*_⌋), up (Δ*y* = ⌊*d*_*avr*_⌋), and down (Δ*y* = −⌊*d*_*avr*_⌋) on ⌊*d*_*avr*_⌋ = 5 pixels, respectively. **n** Output segmentation of two-nuclei cluster based on the perturbed watershed

*Virtual cuts*. In some cases, mostly when cell membrane marker is not present, the initial watershed segmentation step might undersegment the cluster. For such cases, we have developed the virtual cuts method that utilizes non-convex topology of the cluster to identify nuclei centroids that act as seed points for the watershed algorithm. See Methods for implementation details.

### New dataset for segmentation benchmarking

As part of our UNSEG development, we have curated 75 tiff images of tissue sections from eight organs of the extended human gastrointestinal system – appendix, colon, esophagus, gallbladder, liver, pancreas, small intestine, and stomach. The immunofluorescence images were acquired via imaging of formalin-fixed paraffin-embedded (FFPE) tissue sections labeled using Hoechst and fluorescent-dye-conjugated Na^+^K^+^ATPase as respective markers for cell nuclei and membranes (See Methods). The image dimensions are 1000 × 1000. The images were acquired using a 0.95 numerical-aperture objective with 40X magnification, and have a pixel pitch of 0.16 μm/pixel. Our gastrointestinal tissue (GIT) dataset includes images of normal tissues as well as tissues related to chronic inflammation, cancer precursor lesions, and cancer. These images capture a wide range of tissue organization from samples with sparsely located cells to those with very high cell density. Figure 3 shows 12 representative images from the GIT dataset.

**Figure 3.**
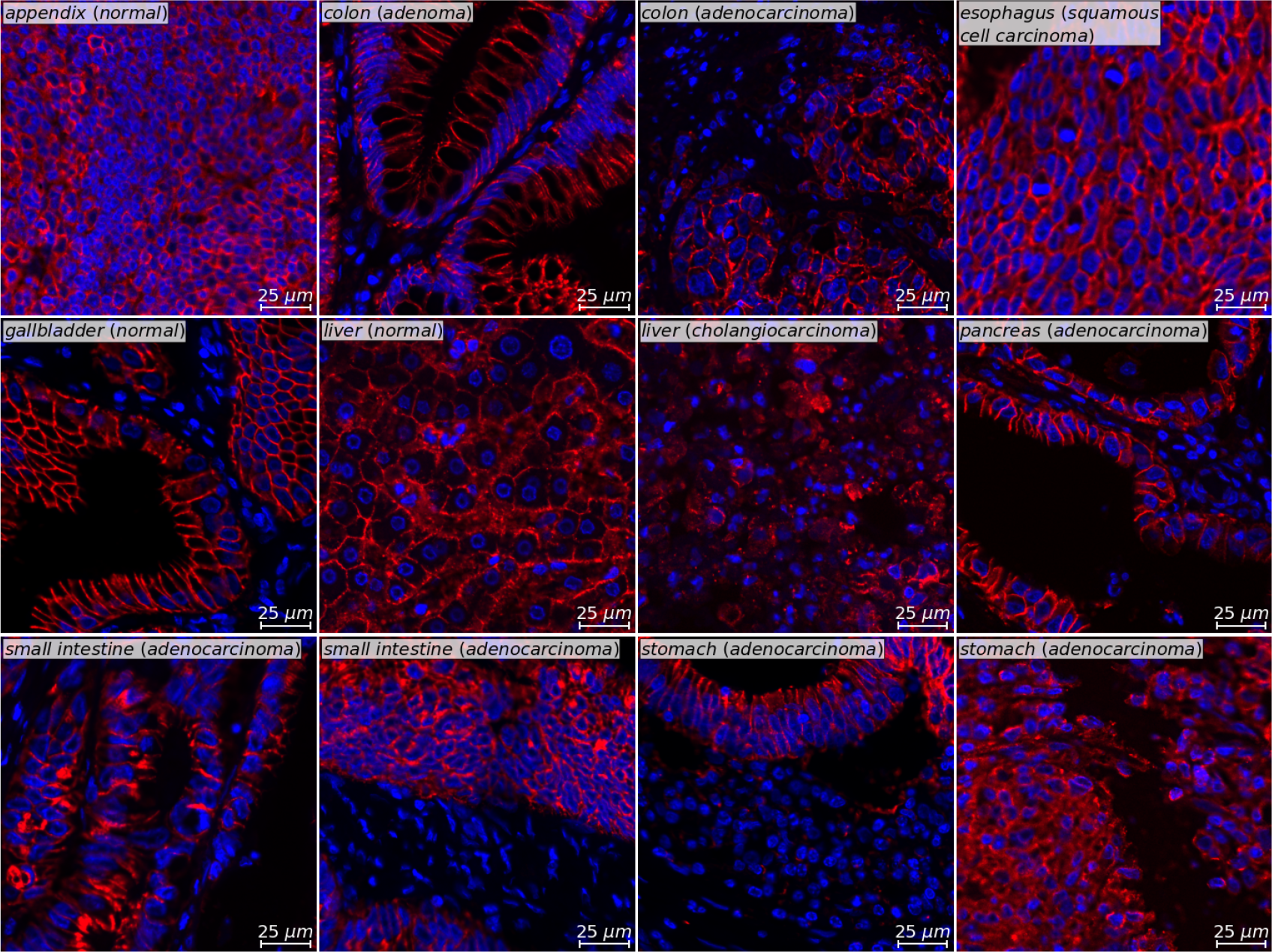
Gastrointestinal tissue (GIT) dataset. Twelve representative tissue images from the GIT dataset drawn from different organs of the human gastrointestinal system with different pathobiology. Blue and red colors respectively indicate nucleus (Hoechst) and cell membrane (Na^+^K^+^ATPase) marker expressions. The dimensions of each image are 1000 × 1000 pixels. The images were acquired using microscope with 0.95 NA, 40X objective and imaging sensor with a pixel pitch of 0.16 μm/pixel.

Expert pathologists independently annotated the 75 images resulting in ground truth with 16201 nuclei and 16217 cells. These annotations were performed manually, without any algorithmic aid, to truly reflect human performance. The detailed description of the dataset is presented in Supplementary Table 1 and Supplementary Figure 1, while the nuclei and cell annotations of 12 representative images are shown in Supplementary Figure 2. To annotate nuclei and cells in the 75 images we developed Cellthon -a Python based graphical user interface for annotating cells and their nuclei in tissue images.

We used the GIT dataset to benchmark UNSEG performance. Moreover, we anticipate that this dataset will also serve as a resource for researchers requiring annotated datasets for future algorithm development and testing^30^.

### UNSEG benchmarking using GIT and publicly available datasets

We used GIT and publicly available datasets to benchmark segmentation performance of UNSEG with respect to Cellpose and Mesmer, the two state-of-the-art DL methods that have consistently demonstrated good performance in segmenting immunofluorescence imaging data particularly in the context of highly multiplexed imaging^25,26^. To perform the comparison with Cellpose, we used Cellpose version 2.1.0. In this version, we chose *nuclei* and *TN2* models from the Cellpose ‘model zoo’ to respectively segment nuclei and cells. Our choice was based on them giving the best segmentation results for the GIT dataset in comparison to all other Cellpose models. We used Cellpose size calibration procedure to estimate the cell diameter for each of the 75 images in our dataset. We also chose Mesmer model, DeepCell 0.12.6, and set the model parameter image_mpp to the pixel pitch in microns per pixel for our imaging dataset. Benchmarking was performed by computing the *F*_1_ score (Eq. 7) as a function of intersection over union (IoU) threshold^46^. The IoU threshold metric quantifies the degree of overlap between algorithm prediction and the annotated ground truth. It is bounded between 0 and 1, with one indicating perfect overlap. By computing the *F*_1_ score over the IoU range, we obtain the *F*_1_ accuracy curve for each method. (See Methods for more details).

Figure 4a shows UNSEG, Cellpose, and Mesmer segmentation results applied to four representative examples from our 75 image GIT dataset. Visual comparison shows similar performance between the different methods. One difference between UNSEG and the other two methods is that, although, UNSEG does implement boundary smoothing, it does not enforce strict shape constraints. As a consequence, the shape of UNSEG-based nucleus and cell segmentation is more irregular, but also more realistic and less synthetic appearing than Cellpose and Mesmer, where the segmented shape and the partitioning of the cell clusters strongly correlates with the nucleus and cell ground truth for the training data.

**Figure 4.**
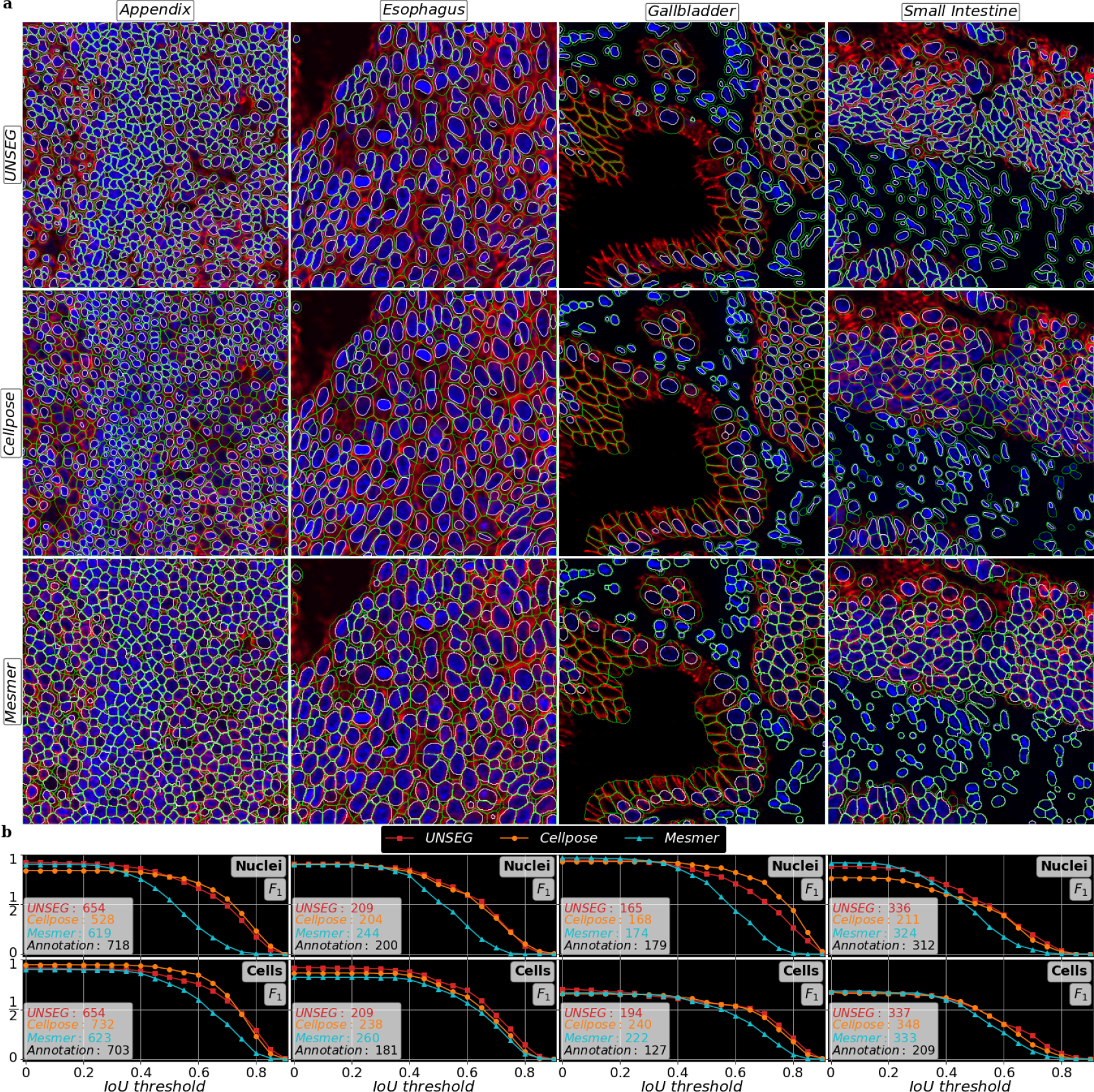
Comparison of UNSEG, Cellpose, and Mesmer on four example images from the GIT dataset. **a** Columns respectively correspond to appendix, esophagus, gallbladder, and small intestine tissue images. Rows show nucleus (white boundary) and cell (green boundary) segmentation results for the four examples using UNSEG, Cellpose and Mesmer, respectively. **b** The two rows respectively show nucleus and cell segmentation accuracy of UNSEG, Cellpose, and Mesmer. Accuracy is measured using number of segmented objects (see inserts) and *F*_1_ score curves plotted as a function of IoU threshold between the segmented and annotated labels for nuclei and cells, respectively.

The *F*_1_ curves for the four examples (Figure 4b) demonstrate that UNSEG performance is similar to that of the DL methods trained on about a million cells. The ground truth annotations for these four examples are shown in Supplementary Figure 2.

The similarity in their performance on the four example images generalizes to the entire GIT dataset. The results are shown in Figure 5. The first row depicts the median *F*_1_ curves corresponding to nucleus and cell segmentation by the three methods. The curves indicate that the three methods have similar segmentation performance. For cell segmentation, the median UNSEG performance is slightly below the other two methods, which is partly due to the conservative nature of UNSEG cell segmentation in resolving cell boundary ambiguity in cases where the tissue section capture partial cell membranes without their respective nuclei. In these cases, UNSEG does not always include their segmentation masks in the final results. (Also see, ‘*F*_1_ score and accuracy’ section below.) Nevertheless, if we look at the pairwise 95% *F*_1_ confidence interval comparison between UNSEG performance, with Cellpose and Mesmer – the second and third rows of Figure 5 respectively – we clearly see their almost complete overlap, indicating their overall similar performance. A more detailed version of Figure 5 is presented in Supplementary Figure 3. We note that we used the same UNSEG parameters to segment all 75 images in the GIT dataset and did not optimize them for every image, despite this ability being a strength of UNSEG and would have boosted its performance. The rationale for eschewing this adjustment was to demonstrate that our probabilistic reinterpretation of the two channel image through a Bayesian lens provides UNSEG with robustness and performance stability, and prevents it from being brittle and requiring continuous adjustment. We additionally note that this is unlike our characterization of Cellpose performance, where we adjusted its size parameter for every image. Therefore, our performance curves are biased towards Cellpose. The UNSEG parameter values we used for GIT dataset are listed in Supplementary Table 2 and discussed in Methods section.

**Figure 5.**
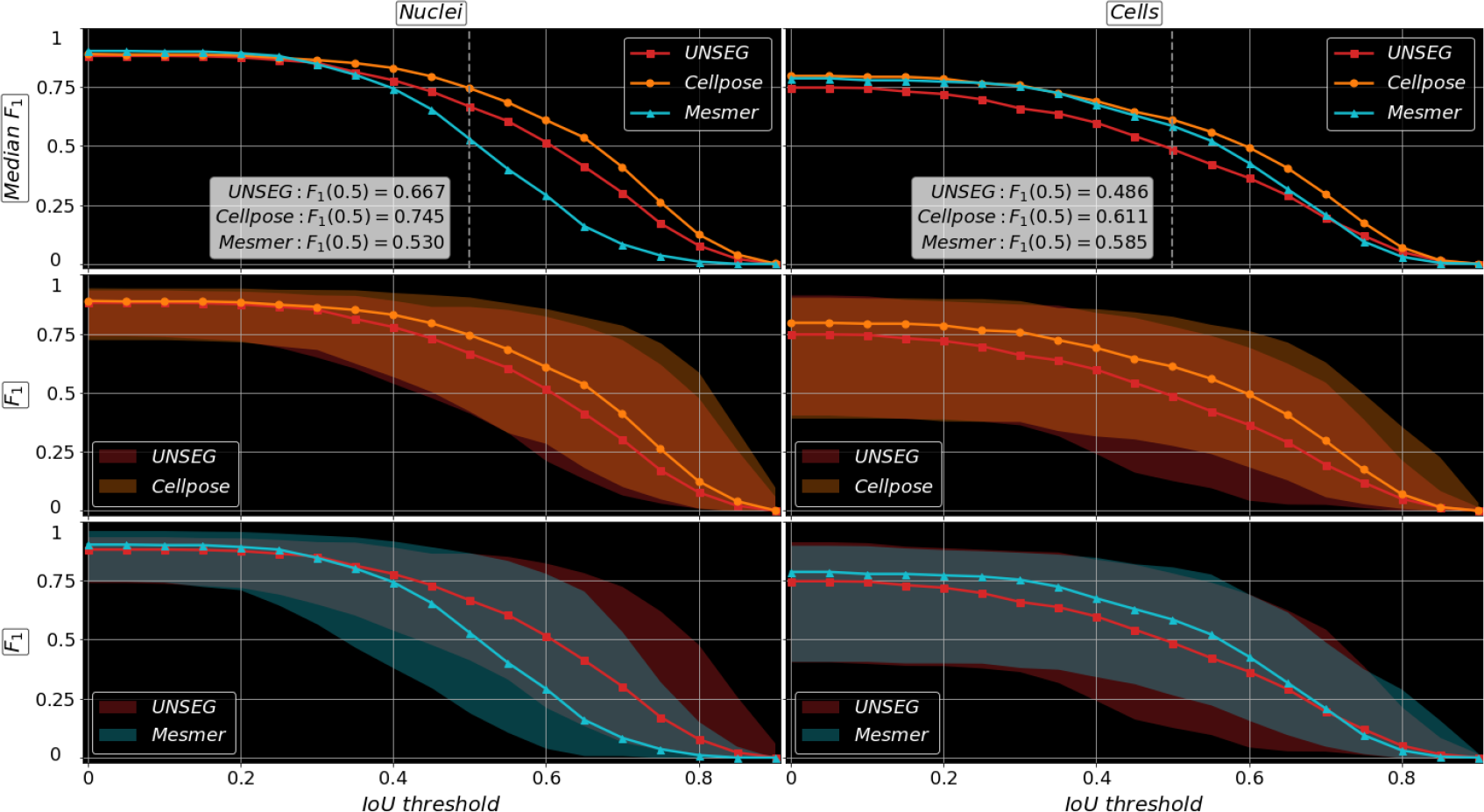
Performance comparison of UNSEG, Cellpose, and Mesmer for the entire GIT dataset. First row compares median *F*_1_ score performance curves for the three methods as a function of IoU threshold for nucleus and cell segmentation of images in the GIT dataset. The insert contains median *F*_1_ score values at the IoU threshold of 0.5 for three algorithms. The second and third rows respectively show pairwise comparison between UNSEG and Cellpose, and UNSEG and Mesmer. The comparison includes median *F*_1_ score curves along with their 95% confidence intervals. Their complete overlap indicates similar performance of all three methods.

Furthermore, we also benchmarked segmentation performance of UNSEG with respect to Cellpose and Mesmer using publicly available, multiplexed imaging tissue datasets acquired using CODEX, Vectra, and Zeiss imaging platforms^47^. Supplementary Figures 4 through 6 respectively show the cell segmentation performance of UNSEG, Cellpose, and Mesmer on CODEX, Vectra and Zeiss datasets. The Codex dataset comprises of ten 400 × 400 images of lymph nodes and tonsils. For our benchmarking we chose CD20 and CD45RO as cell-membrane markers to demonstrate the ability of UNSEG to work with different cell-membrane markers. These images were acquired using an objective with 20X magnification, and imaging sensor with pixel pitch of 0.3774 μm/pixel^47^. Supplementary Figure 4a depicts an example image of lymph node from the CODEX dataset, along with its ground truth cell annotation, the cell segmentation predicted by UNSEG, Cellpose, and Mesmer, and their corresponding *F*_1_ score based performance curves. Due to the high cell density, lymph node samples are typically difficult to segment. This example provides a clear visual and quantitative demonstration of UNSEG performing segmentation on par with Cellpose and Mesmer. Supplementary Figure 4b further shows that the quality UNSEG performance extends to the entire CODEX dataset.

Similarly, Supplementary Figures 5 and 6 compare the performance of UNSEG cell segmentation with that of Cellpose and Mesmer for Vectra and Zeiss datasets^47^ respectively. The Vectra dataset includes 131 tissue images of size 400 × 400 from a range of pathologic diseases that include lung adenocarcinoma, extramammary Paget disease, pancreatic ductal adenocarcinoma, lung small cell carcinoma, colon adenocarcinoma, Hodgkin lymphoma, breast ductal carcinoma, serous ovarian carcinoma, squamous cell carcinoma, Merkel cell carcinoma, and squamous mucosa. The Zeiss dataset consists of nineteen tissue images of size 800 × 800, acquired from tissue sections of cutaneous T-cell lymphoma, pancreatic adenocarcinoma, basal cell carcinoma, and melanoma. Both Vectra and Zeiss datasets were acquired using 20X magnification objectives however pixel pitches of imaging sensors were 0.5 μm/pixel and 0.325 μm/pixel respectively^47^. Although, UNSEG performs stable and high quality segmentation, faithfully capturing cell shapes, its *F*_1_ score based performance is upper bounded by Cellpose and Mesmer. This is partly due to annotated ground truth having a tendency to be over-segmented, which tends to favor Cellpose and Mesmer *F*_1_ scores (also see, ‘*F*_1_ score and accuracy’ section below). We found this to be particularly true for Vectra dataset. For this dataset, it was also difficult to find cell membrane markers that were appropriately imaged across the different images. We, therefore, utilized pan-cytokeratin, a cytoplasmic marker for cell segmentation. Since, UNSEG has been developed for utilizing nucleus and cell membrane marker for unsupervised segmentation, and not nucleus and cytoplasm marker, we did expect reduced performance. However, the quality of UNSEG segmentation remained remarkably robust, despite expected reduction in UNSEG *F*_1_ score values.

### Applicability of UNSEG to different practical scenarios

We also tested UNSEG performance in multiple different practical scenarios.

1. *Weakly expressing cell membrane marker*: We identified a tissue image of human skin with dermatofibrosarcoma acquired from a publicly available CODEX dataset^13^, which is a different dataset from the one discussed above. This image has weakly expressing Na^+^K^+^ATPase as the cell-membrane marker. Hoechst is the nucleus marker. The image size is 1440 × 1440 pixels. It was acquired using an objective with a 20X magnification and a sensor with a pixel pitch of 0.377 μm/pixel. As shown in Supplementary Figure 7, UNSEG demonstrates stable and robust segmentation performance with a weakly expressing membrane marker. As this dataset lacked annotations, we did not compute the *F*_1_ curve but as the figure demonstrates, a visual, qualitative assessment of UNSEG segmentation compares favorably with Cellpose and Mesmer.
2. *Using a non-specific cell-membrane marker to segment cells*: In Supplementary Figure 5, using the Vectra dataset, we demonstrated that UNSEG is robust to using cytoplasmic markers for cell segmentation. To further test the wide applicability of UNSEG, we replaced weakly expressing Na^+^K^+^ATPase with Hyaluronan, which cannot only localize to the cell membrane but also to the cytoplasm and the extra-cellular matrix. We used Hoechst as the nucleus marker. Supplementary Figure 8 shows that UNSEG performs high quality nucleus and cell segmentation, which also compares favorably with generalist methods like Cellpose and Mesmer.
3. *DRAQ5 as the nucleus marker*: We next switched Hoechst with DRAQ5 as the marker for the nucleus, while keeping Hyaluronan as the cell membrane marker. Supplementary Figure 9, show that UNSEG continues to provide high quality segmentation.
4. *Applying UNSEG to multiplexed ion beam imaging (MIBI)*: We also tested UNSEG sub-cellular segmentation performance on nuclei and cells in a placental tissue image acquired using MIBI, an alternative multiplexed imaging technology^6,8^. The image was downloaded from the Human BioMolecular Atlas Program (HuBMAP) database.^48^ The image size is 2048 × 2048, with pixel pitch of 0.391 μm/pixel. Due to lack of clearly identified annotation, Supplementary Figure 10 does not show the *F*_1_ curves, but does provide a visual comparison of UNSEG, Cellpose and Mesmer performance. As before, UNSEG performance continues to be at par with deep learning methods.

### *F*_1_ score and accuracy

*F*_1_ is a well-established score for assessing segmentation accuracy. It simultaneously accounts for the proportion of correctly segmented objects and their pixel-wise matching with ground truth object profiles^46^. However, as we show in Supplementary Figure 11, *F*_1_ score is biased depending on how the estimated segmentation mask deviates from the ground truth. Specifically, *F*_1_ value is higher if the size of the estimated segmentation mask is larger than the ground truth, as compared to when it is smaller. In fact, as shown in Supplementary Figure 11, the former upper bounds the latter. Both Cellpose and Mesmer, on average, have larger cell segmentation mask estimates when compared to UNSEG. This is a contributory factor towards the higher median *F*_1_ scores for Cellpose and Mesmer, even when segmentation results from all three methods are reasonable. Supplementary Figure 4 exemplifies this point. There, even though cell segmentation results from all three methods are reasonable, UNSEG has a slightly lower *F*_1_ curve, due to it being conservative in estimating cell size, as is discussed above in the subsection on UNSEG benchmarking.

### UNSEG characteristics and use case

UNSEG employs an integrated approach to segmenting nuclei and cells that, by design, emphasizes internal consistency between each cell nucleus and its membrane. As a consequence, UNSEG guarantees that no segmented nucleus can be located beyond the boundaries of its cell. This drawback is often found in both Cellpose and Mesmer, where nucleus and cell segmentations are performed independently. Figure 6a depicts a small intestine tissue section illustrating the internal inconsistency in nucleus and cell boundaries estimated by Cellpose and Mesmer for a pair of examples highlighted with dashed boxes. In the case of Cellpose the larger nucleus is located in two cells, while in Mesmer, for region marked as 1, two cells are sharing the same nucleus. For region marked as 2, in the case of Cellpose the nucleus extends beyond the boundary of its cell. UNSEG avoids such discrepancies due to its joint segmentation of nuclei and cells. This joint processing ensures that UNSEG can unambiguously identify the cytoplasmic compartment of cells. The internal consistency among sub-cellular compartments is of particular importance in biological studies where correct sub-cellular localization of signaling pathway components is essential to study intra-cellular signaling. For example, tumor protein P53 can be sequestered in the cytoplasm, or localized in the nucleus depending on DNA damage, and other exogenous and endogenous stresses. However, in unstressed cells, it is expressed at low levels and localizes in both cytoplasm and the nucleus^49^. As another example, histone methyltransferase EZH2 localizes in the nuclei, where it regulates gene expression through its canonical histone lysine methyltransferase activity^50^. Supplementary Figure 12 depicts an example of such a real use case, where UNSEG is used in a multiplexed imaging context to segment cells and their nuclei based on Hoechst and Na^+^K^+^ATPase. The UNSEG-based segmentation is used to localize intra-cellular P53 and EZH2 expression in a region of healthy colon tissue with densely located cells (see Methods). The internal consistency of UNSEG segmentation ensures that the user is correctly able to evaluate P53 expression in the nucleus and the cytoplasm, while ensuring that the canonical activity of EZH2 in the healthy tissue is not associated with the cytoplasm.

**Figure 6.**
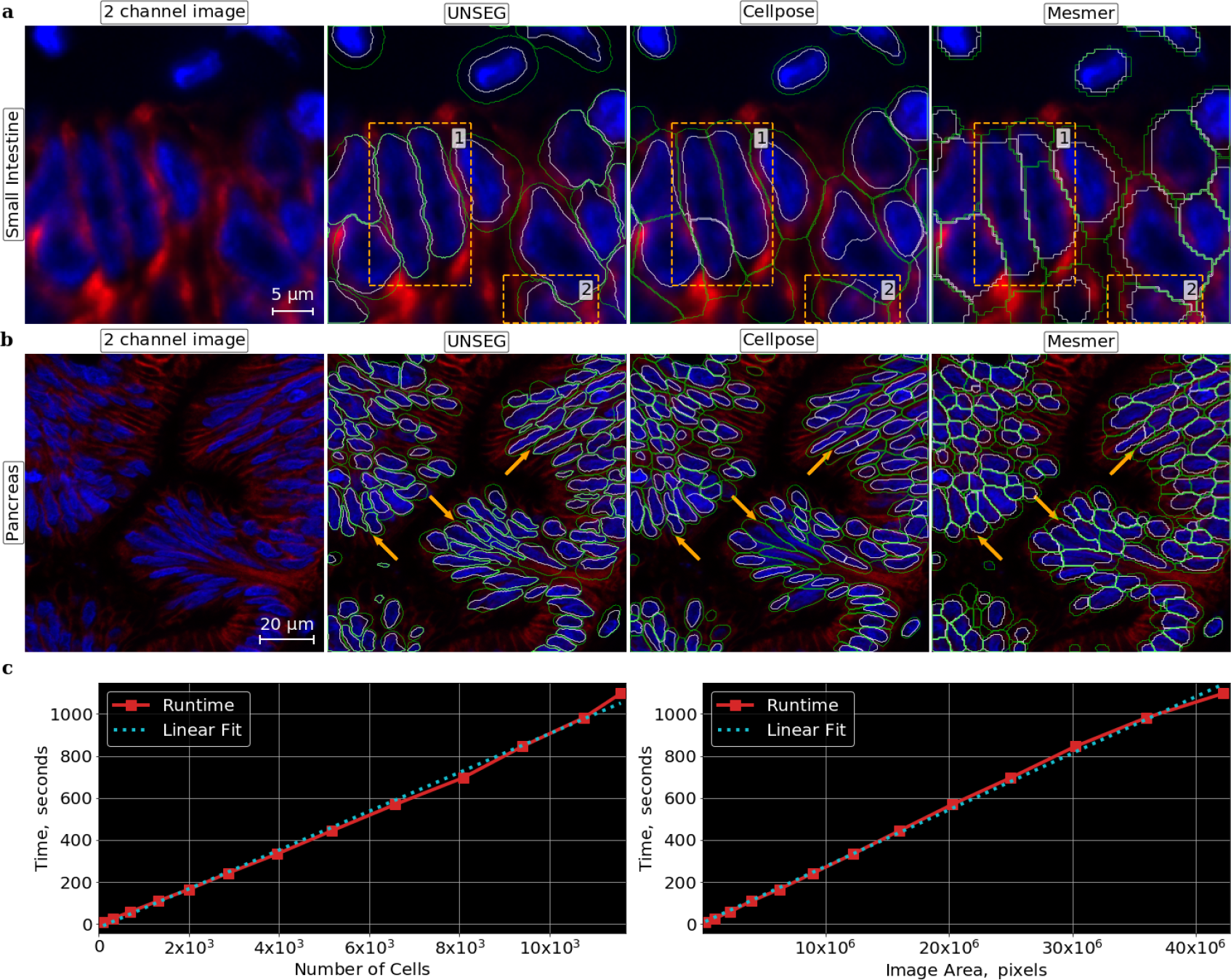
Characteristics of UNSEG method. **a** UNSEG demonstrates internal consistency between nucleus and cell boundaries. The dashed box 1 contains two nuclei, where both Cellpose and Mesmer have mismatch between the boundaries of two cells and their nuclei. The dashed box 2 contains another cell, where Cellpose has mismatch between the nucleus and its cell boundaries. **b** UNSEG is better at capturing complex shapes of nuclei in comparison to Cellpose and Mesmer, as exemplified by the arrows indicating examples of nuclei with complex shapes. **c** Runtime complexity of UNSEG as a function of number of cells and image area, assuming uniform cell distribution for the latter.

As briefly mentioned earlier, UNSEG does not impose a strict shape constraint on the segmented nuclei by allowing them to be locally non-convex. Consequently, in complex tissue sections it is, on average, better at preserving true nucleus shape than Cellpose and Mesmer, which either are usually more rounded, and in regions of the tissue with high cell density, appear like Voronoi partitions of the tissue region. Figure 6b shows an example of pancreas tissue with elongated cells that deviate from round shapes. As can be seen, the ability of UNSEG to combine knowledge of global tissue architecture and local topology, with a relaxed shape constraint allows it to better capture elongated nucleus morphology when compared to Cellpose and Mesmer. This ability is highly relevant in the context of the use case mentioned above, where users, such as cancer biologists are studying the tumor microenvironment that might include a diversity of cell shapes associated with cancer, immune, and stromal cell populations.

Runtime complexity of UNSEG is a function of number of cells and not the image size. Specifically, UNSEG runtime complexity scales approximately linearly with respect to the number of segmented cells in the image. This translates to linear scaling with respect to image area, if the spatial distribution of cells is approximately uniform. However, for sparsely populated images UNSEG run time will be significantly sub-linear. Figure 6c shows linear dependence with respect to the number of segmented cells and the image area, under the assumption of uniform cell distribution. The results were generated using an acquired colon tissue microarray (TMA) spot with approximately uniform cell distribution. The segmentation results for the whole TMA spot are presented in Supplementary Figure 13.

## Discussion

The importance of segmenting cells and their nuclei has gained renewed prominence due to the advent of multiplexed imaging technologies that have significantly enhanced the depth of information that can potentially be extracted from samples in a cell specific manner. However, tissue sections have complex cell organizations and unlike computer vision tasks, segmenting individual cells even by human experts is a difficult challenge, resulting in inter-observer discordance. Such discordance usually grows as the number of cells requiring annotation grows. This, in turn, affects ground truth quality used to train supervised learning models, and is a bottleneck for generating high quality training data. The unsupervised approach provides a complementary paradigm to segmenting complex tissue images without requiring training data. Unsupervised methods are also more adaptable to individual images of varying complexity. However, to the best of our knowledge, until now no method within the unsupervised paradigm had demonstrated performance approaching supervised learning methods, particularly those based on deep learning. As a consequence, none of its advantages were relevant. UNSEG, for the first time, to the best of our knowledge, demonstrates that unsupervised cell and nuclei segmentation can achieve accuracy at par with the current state of the art methods in deep learning. It also introduces the perturbed watershed algorithm, a new standalone algorithm that extends the ability of classical watershed algorithm to correctly segment nucleus clusters. Perturbed watershed is applicable in all cases where the classical version can be used. Finally, like the generalist DL methods, UNSEG is not brittle, and is applicable to a range of tissue types, disease pathologies, nucleus and cell membrane markers, and multiplexed imaging modalities. It achieves accuracy on par with these methods, along with the added benefit of guaranteeing segmentation consistency between a cell and its nucleus, and being faithful to their morphology.

Segmentation fundamentally involves learning features and image representations that help the algorithm identify individual cells and their nuclei. Deep learning models extract these features and representations in a supervised manner. Interestingly, UNSEG performance reveals that there is intrinsic information latent in the topology of cells and nuclei within the tissue context of an individual image that is equivalent to training on one million cells^26^. Importantly, this information can be acquired adaptively for every tissue image. Therefore, it is conceivable to develop adaptive DL methods that perform sub-cellular segmentation of individual unlabeled tissue images adaptively, by leveraging UNSEG as a label generator to initialize internally consistent cell and nucleus labels that a DL method can optimize and improve using self- and semi-supervised learning paradigms. For example, in a self-supervised learning framework UNSEG could be used to optimally initialize joint learning of neural network parameters and *k*-means based segmentation of cells and nuclei^51^. Another application could be in a semi-supervised setting, where a small portion of the image is annotated, while the remaining is unlabeled. Here, UNSEG could be used to provide pseudo-labeling estimate of cell and nucleus segmentation for the unlabeled data, which can then be used to refine the DL model trained on labeled data^52,53^. Finally, UNSEG could be used in the setting of learning with noisy labels, where the UNSEG generated segmentation masks are noisy labels on which robust DL models can be trained^54^.

UNSEG performs sub-cellular segmentation based on nucleus and cell membrane compartment markers. However, its framework does not impose any constraint on the number of markers that can be used. For example, in multi-nucleated cells, UNSEG can be modified to incorporate an additional marker specific to the nuclear membrane to coherently segment multiple overlapping nuclei belonging to the same cell. Supplementary Figure 14 depicts an example of a multi-nucleated cell, with Lamin A/C (shown in green) marking the nucleus membranes. As depicted in this figure, the modification of UNSEG utilizes the specificity of the extra marker to segment the nuclei and associate them with the same cell.

UNSEG is an easy to use method for sub-cellular segmentation of complex tissue images using multiplexed imaging technologies. It only uses well-known and robust python libraries that require minimal setup and is accessible to researchers with varying computational backgrounds. It has two primary parameters (minimal area and convexity threshold; see Methods and Supplementary Table 2.) that can be adjusted by the user to optimize segmentation performance for individual images including relatively large images like shown in Supplementary Figure 13. It is a flexible framework that can be extended to include additional markers to enhance cell segmentation and to extract localized expression of individual markers across the tissue sample. Finally, we reemphasize that unlike segmentation of objects in computer vision based situational awareness tasks, segmenting cells and their nuclei, particularly in the context of tissue samples, often results in subjective ground truth. By being able to capture intrinsic, marker-specific topological structure of cell compartments, UNSEG offers opportunities to further improve current-state-of-the-art deep learning methods. To aid in this task, we have also generated a GIT dataset of 75 tissue images from eight organs of the human gastrointestinal system, along with their corresponding nucleus and cell annotations independently generated by expert pathologists.

## Methods

### Generation of GIT dataset and other images

For GIT dataset, formalin-fixed paraffin-embedded (FFPE) tissue microarray (TMA) slides were obtained from Pantomics (Pantomics, DID381) Tissue TMA samples for Supplementary Figures 12, 13, and 14 were obtained from Department of Pathology at University of Pittsburgh Medical Center Presbyterian Hospital. The slides went through cyclic immunofluorescence antigen retrieval protocol^10^. The corresponding figure slides were stained in cycles with 1:200 dilution of Anti-Sodium Potassium ATPase antibody (Abcam ab198367, clone EP1845Y), 1:100 dilution of P53 antibody (Abcam ab270192, clone SP5), 1:50 dilution of EZH2 antibody (CST 45638, clone D2C9), and 1:100 dilution of LAMIN A*/*C antibody (CST 8617, clone 4C11) overnight at 4°C in the dark, followed by staining with Hoechst 33342 (CST 4082S) for 10 minutes at room temperature in the dark. TMA images were acquired using a 0.95 NA and a 40X objective on a Nikon Ti2E microscope.

Seventy five, 1000 × 1000 high quality regions were identified and extracted from the TMA images and saved as tiff images. Expert pathologists independently annotated these images. The annotations were done using Cellthon, a python based cell annotation graphical user interface (GUI) we created using Tkinter toolkit^55^. Together these 75 images and their cell and nucleus annotations comprise the GIT dataset.

### UNSEG algorithm

#### Input image

The input to our algorithm is a two channel image. An example is illustrated in the “input” panel of Figure 1 and Supplementary Figure 15, as well as in Figure 3 and Supplementary Figure 13. Channel one, depicted in blue, and channel two shown in red, are respectively associated with nucleus and cell-membrane marker expressions. Each channel of the image is independently scaled to 0 and 1, such that *I*_*i*_ : Ω → [0, 1]. Here *I*_*i*_ is the normalized image intensity for *i*-th channel, Ω is the image domain, and *i* = 1, 2 is the indexing representing the two channels.

The algorithm performs nucleus and cell segmentation utilizing a Bayesian framework: the posterior probability estimates of nucleus and cell masks are obtained from their *a priori* and likelihood estimates that UNSEG computes from the normalized two channel image. These posterior estimates are then used to obtain the final nucleus and cell segmentations. UNSEG implements this framework through four processing stages detailed below and illustrated in Figure 1 and Supplementary Figure 15.

#### Processing stage 1: Computing a priori nucleus and cell membrane masks

In Stage 1 we compute *a priori* estimates of image foreground for each channel. The estimates are computed at the global and local scale as described below.

#### *A priori* probability

Each channel, *I*_*i*_(*x, y*), *i* = 1, 2, is first pre-processed using a combination of a Gaussian filter^56^ and multi-level Otsu^56–58^. The standard deviation of the Gaussian filter kernel, *σ* is a parameter of the algorithm that allows the user to control the degree of smoothing. This and other algorithm parameters are summarized in Supplementary Table 2. Our default setting is *σ* = 3. A three-level Otsu is next applied to the smoothed image and the lowest level is selected as the threshold to obtain the initial estimate of the channel foreground.

We use the initial, per-channel foreground estimate to compute the cumulative distribution function (CDF), *ℱ*_*i*_ of *I*_*i*_ using intensity values, *I*_*i*_(*x, y*), of pixels (*x, y*) within this estimate. Two examples of CDFs are presented in Supplementary Figure 15.Using the monotonically non-decreasing property of CDF we map *I*_*i*_ to its cumulative probabilistic representation 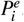, where 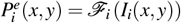.We define 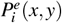 to be the *a priori* probability of the pixel being the nucleus (*i* = 1) or cell membrane (*i* = 2). We note that this definition quantifies the intuition that stronger the marker intensity at a particular pixel, the higher its *a priori* probability. Examples of *a priori* probabilities for nuclei 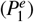 and cell membranes, 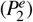 are presented in Figure 1 and Supplementary Figure 15.

#### *A priori* global mask

We compute the *a priori* global mask 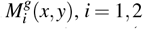 using 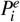 and a simple filter called local mean suppression filter (LMSF) that we have developed. The foreground pixels (*x, y*) where 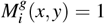 are designed to be a superset of the pixels belonging to the true nucleus (*i* = 1) and cell membrane (*i* = 2) compartments of cells in *I*_*i*_(*x, y*), *i* = 1, 2.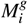, therefore, ensures that no pixels belonging to the cells are missed.

LMSF is designed to identify the valleys (or space) that exist between nuclei (or cell membranes) of closely located cells that nevertheless have some spill over marker expression, and are therefore, difficult to identify as background. We define LMSF as,

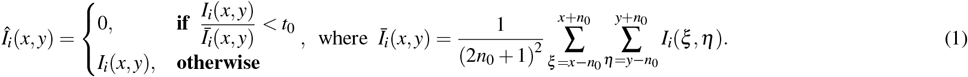

The above definition states that for a given pixel (*x, y*) ∈ Ω, LMSF replaces the original intensity value with 0 only if the ratio of the pixel intensity to the average intensity, computed locally around the pixel neighborhood, is below the threshold parameter *t*_0_.The size of the kernel defining the neighborhood over which the local mean intensity is computed is parameterized by *n*_0_. We set *t*_0_ = 0.5. Consequently, all pixels with intensity value less than half the mean intensity in their respective neighborhoods are replaced with zeros, allowing us to identify valleys between cells. By varying *n*_0_ we can identify valleys and gaps of different widths. UNSEG performs LMSF filtering for *n*_0_ = 5, 10, 20, 40. If Î_*i*_(*x, y*) = 0 for any value of *n*_0_, then the final pixel value is set to 0 and assigned to be background in the global mask,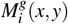. Thus, LMSF allows us to capture valleys of different widths. The values of *n*_0_ are user defined and can be optimized according to complexity of individual images.

We refine the global mask 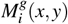 by reassigning those pixels currently in the foreground that have *a priori* probability 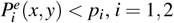 to the background. This refinement is particularly useful for images with highly heterogeneous tissue with varying marker expression. The threshold value *p*_*i*_ should be small and by default is set to 0.01.

An example of *a priori* global mask is presented in Supplementary Figure 15.

#### *A priori* local mask

Complementing 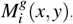,we next compute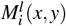, the *a priori* local mask corresponding to image 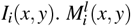 captures the local peculiarities of the compartments – nuclei or cell membranes – associated with their local structure and morphology.

First, *I*_*i*_(*x, y*) is filtered by applying a single iteration of gradient adaptive smoothing (GAS)^45,59^,

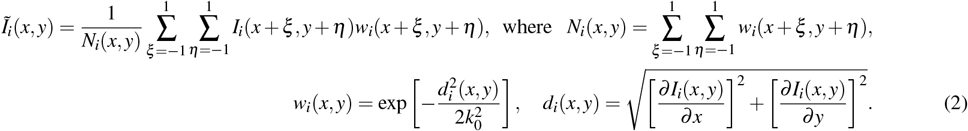

This GAS filtered image, Ĩ_*i*_(*x, y*) smooths the original image, *I*_*i*_(*x, y*), while preserving the local variations within and around cell nuclei and membranes. The local neighborhood is defined via a 3 × 3 kernel, *w*_*i*_, that also performs variation preserving smoothing. Here, variation is quantified via computation of local gradient and the degree of smoothing is controlled by *k*_0_, which is an algorithmic parameter. Its default setting is 1.

To obtain 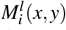, a two-level, local Otsu is applied to Ĩ_*i*_(*x, y*) based on disk kernel whose radius *r*_0_ is an algorithmic parameter. Its default setting is 5 pixels. The Otsu output faithfully captures the local structure but is also noisy, particularly in image regions where no tissue samples are present and the gradients are being computed on the background noise. As 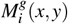 can accurately identify such background, the output of the local Otsu is restricted to where 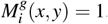, resulting in local foreground mask 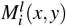.

An example of *a priori* local mask is presented in Supplementary Figure 15.

### Processing stage 2: computing a posteriori nucleus and cell membrane masks

The *a priori* global and local binary masks are computed independently for both channels. As a result, non-negligible probability exists for a pixel to be classified as being both in the nucleus and cell membrane. This is particularly true in tissue regions with crowded cells, or when the nature of the tissue section is such that cell membrane is laying over the nucleus. This processing stage reconciles these overlaps and generates *a posteriori* global and local nucleus and cell membrane masks.

#### Contrast based likelihood function

Human visual perception of cell membranes and nuclei is based on inherent contrast between the two channels. Usually this contrast is visualized via imbuing the individual intensity based channels with colors. Here, we adapt this notion to compute a visual contrast function based on nucleus and cell membrane marker specific expression to quantify the likelihood of pixel belonging to either the nucleus or cell membrane. The first step computes the contrast function for each pixel in the *a priori* local mask as follows,

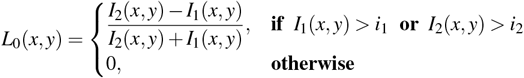

where 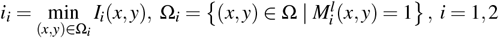. The second step ensures that this function is consistent with the *a priori* global mask for each channel, resulting in the contrast based likelihood function,

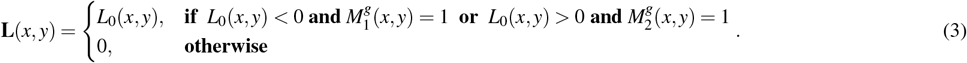

**L**(*x, y*) is bounded between [−1, 1], with the contrast of −1 indicating the strong likelihood that the pixel (*x, y*) belongs to the nucleus, while 1 indicating the pixel most likely belongs to the cell membrane. Two examples of likelihood function are presented in Figure 1 and Supplementary Figure 15.

#### *A posteriori* global mask

We combine the *a priori* probability with the contrast based likelihood function to compute the *a posteriori* global mask **M**^*g*^(*x, y*), such that **M**^*g*^ : Ω → {0, 1, 2}, where the labels 0, 1, and 2 correspond to the background, nuclei, and cell membranes, respectively. However, before performing this combination, we enhance 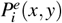 as follows,

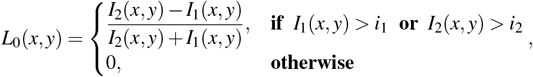

where *i* = 1, 2. This enhancement, saturates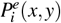 – that is, sets 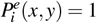– where the *a priori* local mask is 1. It ensures graceful performance of our algorithm in the global context, when computing *a posteriori* global mask **M** (*x, y*). We then compute the *a posteriori* global probability 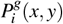, via 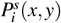-weighted convex combination of the likelihood and *a priori* belief,

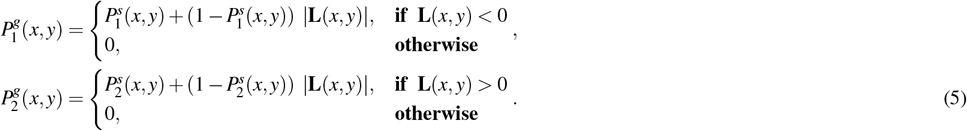

The final posterior global mask is obtained by either applying *k*-*means* clustering, with *k* = 3, or *arg max* operation^45^ on 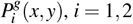. (Eq. 5) to compute **M**^*g*^(*x, y*). The default setting is *arg max*. We note that *k*-*means* (or *arg max*) is performed under the constraint that pixel (*x, y*) ∈ Ω is assigned to the common background if both global probabilities have zeros values,i.e.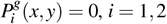. Examples of the *a posteriori* global mask are presented in Figure 1 and Supplementary Figure 15.

#### *A posteriori* local mask

We define the *a posteriori* local mask, **M**^*l*^ : Ω → {0, 1, 2}, simply by restricting the *a priori* probability 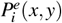 to the local mask 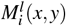,

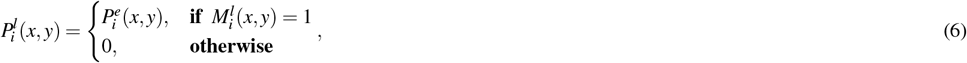

where *i* = 1, 2. This restriction allows us to optimally capture the local *a posteriori* structure of the nuclei and cell membranes in a self-consistent manner.

Similar to computing the *a posteriori* global mask, we either apply *k*-*means* clustering or *arg max* (default setting) operation on 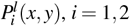 (Eq. 6) to obtain the *a posteriori* local mask **M**^*l*^(*x, y*). As mentioned above for the *a posteriori* global mask, the same constraint for the common background is also applied here. Examples are presented in Figure 1 and Supplementary Figure 15.

#### Processing stage 3: nucleus segmentation

The *a posteriori* global and local masks provide a semantic segmentation of image pixels comprising the tissue into nuclei and cell membranes. This, and the following processing stages are designed to obtain every instance of individual nucleus and its cell from the semantic segmentation of the tissue. Specifically, in this stage, we first segment all nuclei, and use them as a basis to identify their cells in the next stage. These steps ensure that the nucleus and cell segmentations are internally consistent with the latter always bounding the former.

To segment nuclei we process the *a posteriori* global mask for the nuclei, 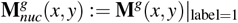 with help from the *a posteriori* local mask for the cell membrane, 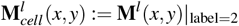 .Particular examples of these two masks are presented in Supplementary Figure 15.

#### Convexity analysis

Nucleus segmentation begins with convex analysis of every connected component of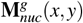. As a part of this analysis, we compute area and the steepest concave point (SCP)^37^ of every component. SCP is a boundary point of the component with the largest deviation from its convex hull. The area parameter allows us to filter out exceedingly small objects that are not nuclei, while SCP helps us determine if the component is nucleus cluster (NC) or not. The component is kept for further analysis only if the area of the component exceeds *a*_0_. Otherwise it is removed. Each component that passes the area threshold, is either classified as an NC or non-NC depending on whether SCP is above or below the threshold *d*_0_. Both *a*_0_ – default set to 20 pixels – and *d*_0_ – default value is 4 pixels – are the primary algorithm parameters (Supplementary Table 2). The non-NC components are statistically analyzed to obtain the initial segmentation for all individual nuclei, along with a small component (SC) list comprising of small convex objects that we are less confident about being nuclei.

Convexity analysis of 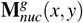, is illustrated in Supplementary Figure 15.

#### Perturbed watershed and virtual cuts

We process the NC components using perturbed watershed (PW) and virtual cut (VC) algorithms that we have developed. Their goal is to partition the NC into individual nuclei.

PW steps are illustrated in Figure 2. Briefly, the NC component mask (Figure 2d) is first modified by 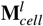 (Figure 2e).Specifically, cuts are introduced in the NC component mask where the local cell membrane is indicated in the 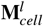 spatially-corresponding to the NC component (Figure 2f). We next apply distance transform (DT) on the modified NC component and use the resulting DT image (Figure 2g) to compute *d*_*avr*_ – the average of all non-zero DT values in the DT image. *d*_*avr*_ is used to threshold the distance transform to identify *n* sub-regions with large DT values indicative of interior of the sub-regions – putative nuclei – making up the NC splitting (Figure 2h). Within every sub-region we identify a pixel with the maximal distance transform value as the watershed seed point (marker) for that sub-region. We perform watershed segmentation of NC based on these *n* seed points to obtain our initial estimate of the nuclei comprising the NC (Figure 2i). If these estimates are correct, then perturbing the markers does not affect segmentation of the NC. However, if the estimates are incorrect, then sub-region estimates are not stable on perturbation. We exploit this perturbation-based stability to identify the correct segmentation of the NC. Specifically, we perturb the marker location and recompute the watershed based segmentation. The perturbations are implemented by shifting each watershed marker location sequentially in the horizontal and vertical directions by ±⌊*d*_*avr*_⌋, resulting in four perturbations: (*x* _*j*_ ± ⌊*d*_*avr*_⌋, *y* _*j*_) and (*x* _*j*_, *y* _*j*_ ± ⌊*d*_*avr*_⌋) with *j* = 1, …, *n* (Figures 2j−2m). Here, ⌊·⌋ stands for the floor function. If during any of the four scenarios, the size of any of the *n* putative nuclei collapses to a point object with an area size bounded to a few pixels (Figures 2j, 2l, and 2m), we deem them as unstable and remove their corresponding seed points from the list of *n* seed points, and recompute the watershed based segmentation with the remaining seed points (Figures 2n). If the segmentation results remain stable for all four shifts, then the estimate is considered correct. To ensure that each of the segmented sub-regions are indeed nuclei and not smaller NCs, we recursively perform convexity analysis and PW on each sub-region. An example of this recursion is illustrated in Supplementary Figure 16.

The above recursive segmentation of an NC can sometimes result in a specific pathological situation, where the convex analysis identifies a sub-region as an NC, but PW does not segment it into sub-regions. For this specific scenario, we have developed the virtual cuts (VC) approach, where a virtual cut is defined through the SCP of the NC component mask to identify virtual sub-regions. We use ‘virtual’ to emphasize that this cut and the resulting sub-regions are only used to identify their respective watershed seed points based on which we perform the actual segmentation. The hypothesis driving the VC method is based on the idea of PW method: although the locations of the respective watershed markers identified using virtual cuts might not exactly coincide with their true locations, they do represent a perturbed version of the true location. Thus, they yield stable and accurate segmentation into the two sub-regions. These sub-regions follow the same recursive logic of the PW method detailed above. VC method is illustrated in Supplementary Figure 15.

Finally, we process the small components in the SC list in a context dependent manner, with small isolated SCs included in the final nucleus segmentation result. Multiple examples of nucleus segmentation are presented in Figures 1, 4, and 6 as well as in Supplementary Figures 7-10, 12, 13, and 15, where the contours of nuclei are outlined in white.

### Processing stage 4: cell segmentation

We segment cells via joint use of *a posteriori* global mask for the cell membranes 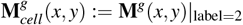 and the segmented nuclei.

We begin by initializing the segmented cell mask as the segmented nucleus mask. The cell mask is then expanded till its boundary coincides with that of the closest cell membrane around it. It is possible that the cell membrane marker used for cell segmentation is not expressed by all cells. Therefore, for cells without any cell-membrane marker expression, the nucleus mask is morphologically dilated a small amount *u*_0_ (1-10 pixels) to obtain an estimate of the cell membrane. *u*_0_ with its 9 pixels default value is one more algorithm parameter (Supplementary Table 2). In the opposite scenario, where due to the nature of the tissue section, a cell is present with a membrane but without a nucleus, we utilize 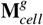. Specifically, the skeleton of 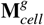 is computed and subtracted from 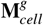 itself. This operation naturally reveals the cell membrane contour within 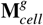 which we identify via computing the Euler number of its connected component. When the Euler number is zero and the area of the connected component exceeds half of the average area of nuclei, the connected component is identified as the segmented cell. Examples of cell segmentation are presented in Figures 1, 4, and 6 as well as in Supplementary Figures 4-10, 12, 13, and 15, where the contours of the segmented cells are outlined in green.

### Performance evaluation

To evaluate UNSEG performance and compare it with Cellpose and Mesmer results, we used the *F*_1_ score (or Dice coefficient) as the accuracy metric^46^. To compute the *F*_1_ score, we first estimated the true positive (*TP*), false positive (*FP*) and false negative (*FN*) values by comparing the predicted segmentation with the expert annotated ground truth and using intersection over union (IoU) as the threshold value^46^. The IoU threshold, ranging from 0 to 1, indicates how much of an overlap between the predicted segmentation and ground truth is considered a match, which is then used to estimate number of *TP, FP*, and *FN* segmented objects. The *F*_1_ score is then given by

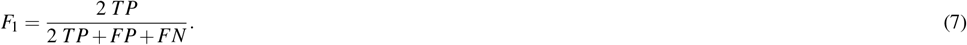

Varying the IoU threshold from 0 to 1, gives us the corresponding *F*_1_ curve as a function of the IoU threshold.

## Data availability

The annotated dataset consisting of 75 fluorescence images of normal and pathologically changed tissue sections of eight organs of the human gastrointestinal system will be made publicly available.

## Code availability

The Python implementation of the UNSEG is available at https://github.com/uttamLab/UNSEG.git. Both Linux and Windows versions of Cellthon are publicly available at https://github.com/uttamLab/cellthon.git.

## Author contributions

S.U conceived the idea. B.K and S.U. designed the overall UNSEG algorithm and planned the key steps. B.K. wrote the UNSEG code and performed the analysis. R.R performed the immunofluorescence labeling and acquired the imaging data. B.R, B.K, R.R, and S.U. identified the 75 images for GIT dataset. P.B., P.S.G, A.S.S, and A.S. provided expert annotation of the GIT dataset images. B.J.L, L.S. R.M.B, R.E.B, J.Y., L.Z., B.D., R.E.S. and A.S. helped in tissue sample acquisition, assay optimization, and data generation. B.K. and S.U wrote the manuscript. All authors reviewed and edited the manuscript before submission.

## Funding

This project was supported in part by the National Institutes of Health through Grant Number UL1TR001857.

## Notes

### Competing Interest Statement

The authors have declared no competing interest.

### Summary of Updates

This revised extends testing of UNSEG to a wide variety publicly available datasets and practical scenarios. It also discusses the fraught nature of quantifying segmentation accuracy in the context of complex tissue samples. In this context it also discusses bias inherent in quantification of segmentation accuracy based on F1 score. Finally, the manuscript includes a more streamlined and updated presentation of results.

## References

1. Wählby, C., Erlandsson, F., Bengtsson, E. & Zetterberg, A. Sequential immunofluorescence staining and image analysis for detection of large numbers of antigens in individual cell nuclei. Cytom. The J. Int. Soc. for Anal. Cytol. 47, 32–41 (2002).

2. Schubert, W. et al. Analyzing proteome topology and function by automated multidimensional fluorescence microscopy. Nat. Biotechnol. 24, 1270–1278 (2006).

3. Zrazhevskiy, P. & Gao, X. Quantum dot imaging platform for single-cell molecular profiling. Nat. Commun. 4, 1619 (2013).

4. Gerdes, M. J. et al. Highly multiplexed single-cell analysis of formalin-fixed, paraffin-embedded cancer tissue. Proc. Natl. Acad. Sci. 110, 11982–11987 (2013).

5. Giesen, C. et al. Highly multiplexed imaging of tumor tissues with subcellular resolution by mass cytometry. Nat. methods 11, 417–422 (2014).

6. Angelo, M. et al. Multiplexed ion beam imaging of human breast tumors. Nat. medicine 20, 436–442 (2014).

7. Lin, J.-R., Fallahi-Sichani, M. & Sorger, P. K. Highly multiplexed imaging of single cells using a high-throughput cyclic immunofluorescence method. Nat. Commun. 6, 8390 (2015).

8. Keren, L. et al. A structured tumor-immune microenvironment in triple negative breast cancer revealed by multiplexed ion beam imaging. Cell 174, 1373–1387 (2018).

9. Gut, G., Herrmann, M. D. & Pelkmans, L. Multiplexed protein maps link subcellular organization to cellular states. Science 361, eaar7042 (2018).

10. Lin, J.-R. et al. Highly multiplexed immunofluorescence imaging of human tissues and tumors using t-cycif and conventional optical microscopes. elife 7, e31657 (2018).

11. Goltsev, Y. et al. Deep profiling of mouse splenic architecture with codex multiplexed imaging. Cell 174, 968–981 (2018).

12. Radtke, A. J. et al. Ibex: A versatile multiplex optical imaging approach for deep phenotyping and spatial analysis of cells in complex tissues. Proc. Natl. Acad. Sci. 117, 33455–33465 (2020).

13. Schürch, C. M. et al. Coordinated cellular neighborhoods orchestrate antitumoral immunity at the colorectal cancer invasive front. Cell 182, 1341–1359 (2020).

14. Phillips, D. et al. Highly multiplexed phenotyping of immunoregulatory proteins in the tumor microenvironment by codex tissue imaging. Front. Immunol. 12, 687673 (2021).

15. Kinkhabwala, A. et al. Macsima imaging cyclic staining (mics) technology reveals combinatorial target pairs for car t cell treatment of solid tumors. Sci. reports 12, 1911 (2022).

16. Hickey, J. W. et al. Spatial mapping of protein composition and tissue organization: a primer for multiplexed antibody-based imaging. Nat. Methods 19, 284 – 295 (2022).

17. Krizhevsky, A., Sutskever, I. & Hinton, G. E. Imagenet classification with deep convolutional neural networks. Adv. neural information processing systems 25 (2012).

18. Goodfellow, I., Bengio, Y. & Courville, A. Deep learning (MIT press, 2016).

19. Ronneberger, O., Fischer, P. & Brox, T. U-net: Convolutional networks for biomedical image segmentation. In Medical image computing and computer-assisted intervention–MICCAI 2015: 18th international conference, Munich, Germany, October 5-9, 2015, proceedings, part III 18, 234–241 (Springer, 2015).

20. Van Valen, D. A. et al. Deep learning automates the quantitative analysis of individual cells in live-cell imaging experiments. PLoS computational biology 12, e1005177 (2016).

21. Al-Kofahi, Y., Zaltsman, A. B., Graves, R., Marshall, W. A. & Rusu, M. A deep learning-based algorithm for 2-d cell segmentation in microscopy images. BMC Bioinforma. 19 (2018).

22. Schmidt, U., Weigert, M., Broaddus, C. & Myers, E. W. Cell detection with star-convex polygons. In International Conference on Medical Image Computing and Computer-Assisted Intervention (2018).

23. Yang, L. et al. Nuset: A deep learning tool for reliably separating and analyzing crowded cells. PLoS Comput. Biol. 16 (2019).

24. Hollandi, R. et al. nucleaizer: a parameter-free deep learning framework for nucleus segmentation using image style transfer. Cell Syst. 10, 453–458 (2020).

25. Stringer, C., Wang, T., Michaelos, M. & Pachitariu, M. Cellpose: a generalist algorithm for cellular segmentation. Nat. Methods 18, 100–106 (2020).

26. Greenwald, N. F. et al. Whole-cell segmentation of tissue images with human-level performance using large-scale data annotation and deep learning. Nat. biotechnology 40, 555 – 565 (2021).

27. Yapp, C. et al. Unmicst: Deep learning with real augmentation for robust segmentation of highly multiplexed images of human tissues. Commun. Biol. 5, 1263 (2022).

28. Lee, M. Y. et al. Cellseg: a robust, pre-trained nucleus segmentation and pixel quantification software for highly multiplexed fluorescence images. BMC bioinformatics 23, 46 (2022).

29. Blazek, P. J. & Lin, M. M. Explainable neural networks that simulate reasoning. Nat. Comput. Sci. 1, 607–618 (2021).

30. Stringer, C. & Pachitariu, M. Cellpose 2.0: how to train your own model. Nat. Methods 19, 1634 – 1641 (2022).

31. Lin, G. et al. A hybrid 3d watershed algorithm incorporating gradient cues and object models for automatic segmentation of nuclei in confocal image stacks. Cytom. Part A 56A (2003).

32. Lin, G. et al. Hierarchical, model-based merging of multiple fragments for improved three-dimensional segmentation of nuclei. Cytom. Part A 63A (2005).

33. Li, G. et al. 3d cell nuclei segmentation based on gradient flow tracking. BMC Cell Biol. 8, 40, 1–10 (2007).

34. Li, G. et al. Segmentation of touching cell nuclei using gradient flow tracking. J. Microsc. 231, 47–58 (2008).

35. Coelho, L. P., Shariff, A. & Murphy, R. F. Nuclear segmentation in microscope cell images: A hand-segmented dataset and comparison of algorithms. 2009 IEEE Int. Symp. on Biomed. Imaging: From Nano to Macro 518–521 (2009).

36. Lou, X., Koethe, U., Wittbrodt, J. & Hamprecht, F. A. Learning to segment dense cell nuclei with shape prior. In 2012 IEEE Conference on Computer Vision and Pattern Recognition, 1012–1018 (IEEE, 2012).

37. Qi, J. et al. Drosophila eye nuclei segmentation based on graph cut and convex shape prior. In 2013 IEEE International Conference on Image Processing, 670–674 (IEEE, 2013).

38. Stoeger, T., Battich, N., Herrmann, M. D., Yakimovich, Y. & Pelkmans, L. Computer vision for image-based transcriptomics. Methods 85, 44–53 (2015).

39. Isack, H. N., Gorelick, L., Ng, K., Veksler, O. & Boykov, Y. K-convexity shape priors for segmentation. In European Conference on Computer Vision (2018).

40. Kostrykin, L., Schnörr, C. & Rohr, K. Globally optimal segmentation of cell nuclei in fluorescence microscopy images using shape and intensity information. Med. image analysis 58, 101536 (2019).

41. Winter, M. R. et al. Separating touching cells using pixel replicated elliptical shape models. IEEE Transactions on Med. Imaging 38, 883–893 (2019).

42. Xie, X. et al. Instance-aware self-supervised learning for nuclei segmentation. In Medical Image Computing and Computer Assisted Intervention–MICCAI 2020: 23rd International Conference, Lima, Peru, October 4–8, 2020, Proceedings, Part V 23, 341–350 (Springer, 2020).

43. Wolf, S., Lalit, M., McDole, K. & Funke, J. Unsupervised learning of object-centric embeddings for cell instance segmentation in microscopy images. In Proceedings of the IEEE/CVF International Conference on Computer Vision, 21263–21272 (2023).

44. Gonzalez, R. C. & Woods, R. E. Digital image processing (Pearson, 2018).

45. Toennies, K. D. Guide to medical image analysis (Springer, 2017).

46. Caicedo, J. C. et al. Evaluation of deep learning strategies for nucleus segmentation in fluorescence images. Cytom. Part A 95, 952–965 (2019).

47. Aleynick, N. et al. Cross-platform dataset of multiplex fluorescent cellular object image annotations. Sci. Data 10 (2023).

48. Human biomolecular atlas program HBM439.HFGX.695. https://portal.hubmapconsortium.org/browse/dataset/54eec389e909636837ccb11958035552.2023-03-09.

49. Maki, C. G. p53 Localization, 117–126 (Springer US, Boston, MA, 2010).

50. Huang, J. et al. The noncanonical role of ezh2 in cancer. Cancer science 112, 1376–1382 (2021).

51. Caron, M., Bojanowski, P., Joulin, A. & Douze, M. Deep clustering for unsupervised learning of visual features. In Proceedings of the European conference on computer vision (ECCV), 132–149 (2018).

52. Lucas, T., Weinzaepfel, P. & Rogez, G. Barely-supervised learning: Semi-supervised learning with very few labeled images. In Proceedings of the AAAI Conference on Artificial Intelligence, vol. 36, 1881–1889 (2022).

53. Arazo, E., Ortego, D., Albert, P., O’Connor, N. E. & McGuinness, K. Pseudo-labeling and confirmation bias in deep semi-supervised learning. In 2020 International joint conference on neural networks (IJCNN), 1–8 (IEEE, 2020).

54. Zheltonozhskii, E., Baskin, C., Mendelson, A., Bronstein, A. M. & Litany, O. Contrast to divide: Self-supervised pretraining for learning with noisy labels. In Proceedings of the IEEE/CVF Winter Conference on Applications of Computer Vision, 1657–1667 (2022).

55. Van Rossum, G. The Python Library Reference, release 3.8.2 (Python Software Foundation, 2020).

56. Chityala, R. & Pudipeddi, S. Image processing and acquisition using Python (Chapman and Hall/CRC, 2020).

57. Otsu, N. A threshold selection method from gray level histograms. IEEE Transactions on Syst. Man, Cybern. 9, 62–66 (1979).

58. Liao, P.-S.Chen, T.-S. & Chung, P. C. A fast algorithm for multilevel thresholding. J. Inf. Sci. Eng. 17, 713–727 (2001).

59. Saint-Marc, P.Chen, J.-S. & Medioni, G. Adaptive smoothing: A general tool for early vision. IEEE Transactions on Pattern Analysis & Mach. Intell. 13, 514–529 (1991).

