## Supplementary data and information for "UNSEG: unsupervised segmentation of cells and their nuclei in complex tissue samples"

| **Type of Tissue** | **Number of Annotated Images** | **Number of Annotated Nuclei** | **Number of Annotated Cells** |
|---|---|---|---|
| Appendix | 4 | 2087 | 2081 |
| -normal | 4 | | |
| Colon | 18 | 4135 | 5060 |
| -normal | 3 | | |
| -chronic colitis | 3 | | |
| -adenoma | 3 | | |
| -adenocarcinoma | 9 | | |
| Esophagus | 9 | 1712 | 1382 |
| -normal | 2 | | |
| -chronic esophagitis | 2 | | |
| -squamous cell carcinoma | 5 | | |
| Gallbladder | 4 | 736 | 737 |
| -normal | 1 | | |
| -chronic cholecystitis | 1 | | |
| -adenocarcinoma | 2 | | |
| Liver | 11 | 1709 | 2316 |
| -normal | 2 | | |
| -cholangiocarcinoma | 2 | | |
| -chronic viral hepatitis | 1 | | |
| -viral subfulminant hepatitis | 1 | | |
| -viral hepatitis related cirrhosis | 1 | | |
| -hepatocellular carcinoma | 4 | | |
| Pancreas | 5 | 970 | 724 |
| -normal | 2 | | |
| -adenocarcinoma | 3 | | |
| Small Intestine | 13 | 2491 | 1994 |
| -normal | 5 | | |
| -chronic enteritis | 3 | | |
| -granuloma | 1 | | |
| -adenocarcinoma | 4 | | |
| Stomach | 11 | 2361 | 1923 |
| -normal gastric mucosa | 2 | | |
| -gastrointestinal stromal tumor | 1 | | |
| -adenocarcinoma | 8 | | |

Supplementary Table 1: Composition of gastrointestinal tissue (GIT) dataset. It includes 75 images of tissue samples from eight different organs of the extended human gastrointestinal system, with different pathobiology conditions.

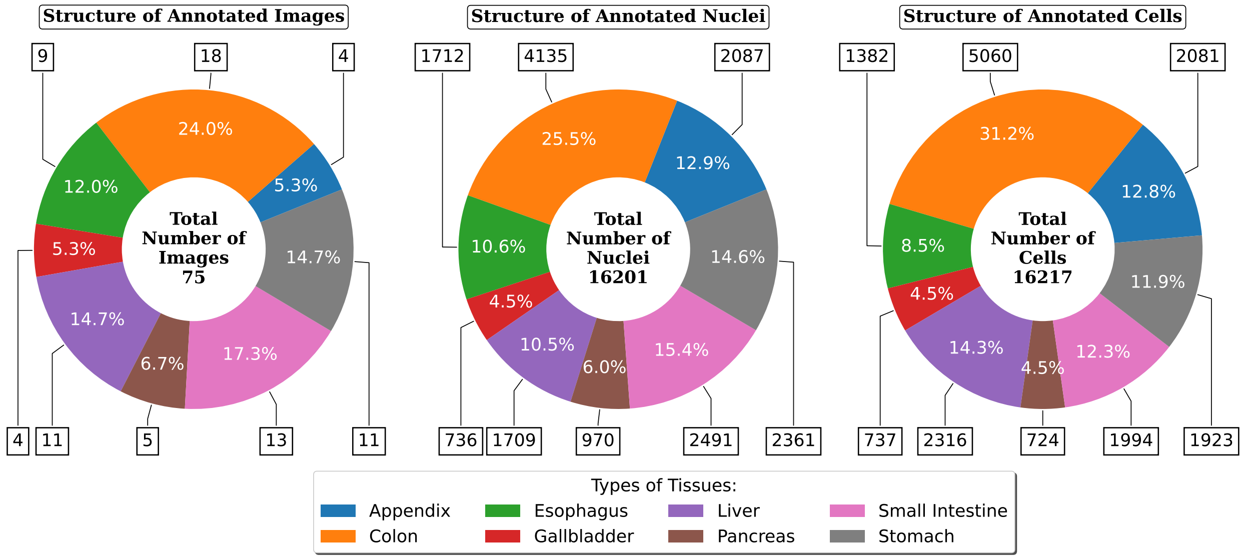


Supplementary Figure 1: Structure of the GIT dataset indicating the percentage of 75 images belonging to each of the eight organs. The number and percentage of expert-annotated nuclei and cells in each of the organs is also indicated.

| Symbol | Description | Python variable name | Default Value | Adjustment Range |
|---|---|---|---|---|
| $a_0$ | The minimal possible area of a nucleus in pixels. | `area_threshold` | 20 | $1 - 200$ |
| $d_0$ | The largest allowable deviation (in pixels) of a boundary point from its convex hull for a convex component. | `convexity_threshold` | 4 | $1 - 10$ |
| $u_0$ | The nucleus mask is morphologically dilated by this amount for cells without cell membrane marker expression. | `dilation_radius` | 9 | $1 - 10$ |
| – | Defines whether a cell membrane marker is present in a cluster of nuclei. | `cell_marker_threshold` | 25 | – |
| – | Defines geometrical distance transform (DT) or gradient distance transform (GDT) in the Virtual Cuts. | `dist_tr` | `'GDT'` | `'DT'` or `'GDT'` |
| $\sigma$ | The standard deviation of the Gaussian filter. | `sigma0` | 3 | – |
| $k_0$ | The degree of smoothing of the gradient adaptive smoothing filter. | `k0` | 1 | – |
| $r_0$ | The disk kernel radius of the local Otsu method. | `r0` | 5 | $1 - 10$ |
| $p_i$ | The background *a priori* probability thresholds for every of two channels. | `pct` | $[0.01, 0.01]$ | – |
| $n_0$ | The list of kernel sizes in the local mean suppression filter. | `nk` | $[5, 10, 20, 40]$ | $1 - 100$ |
| $t_0$ | The intensity threshold in the local mean suppression filter. | `t0` | 0.5 | – |
| – | Defines the algorithm to compute a posteriori local and global masks. | `ternary_met` | `'Argmax'` | `'Argmax'` or `'Kmeans'` |
| – | The area threshold for a cell without a nucleus. | `area_ratio_threshold` | 0.65 | – |
| – | Plots a priori probabilities, a posteriori local and global masks, contrast based likelihood function, and nuclei and cell segmentations. | `visualization` | `False` | `True` or `False` |

Supplementary Table 2: UNSEG parameters with their default values and adjustment range. The primary UNSEG parameters are $a_0$ and $d_0$. If the basic UNSEG algorithm needs to be optimized for an individual image or a set of images then these parameters should be adjusted. The remaining parameters provide only marginal change in segmentation quality, but can be used to fine-tune the performance. The default values and reasonable ranges for their adjustment are also indicated.

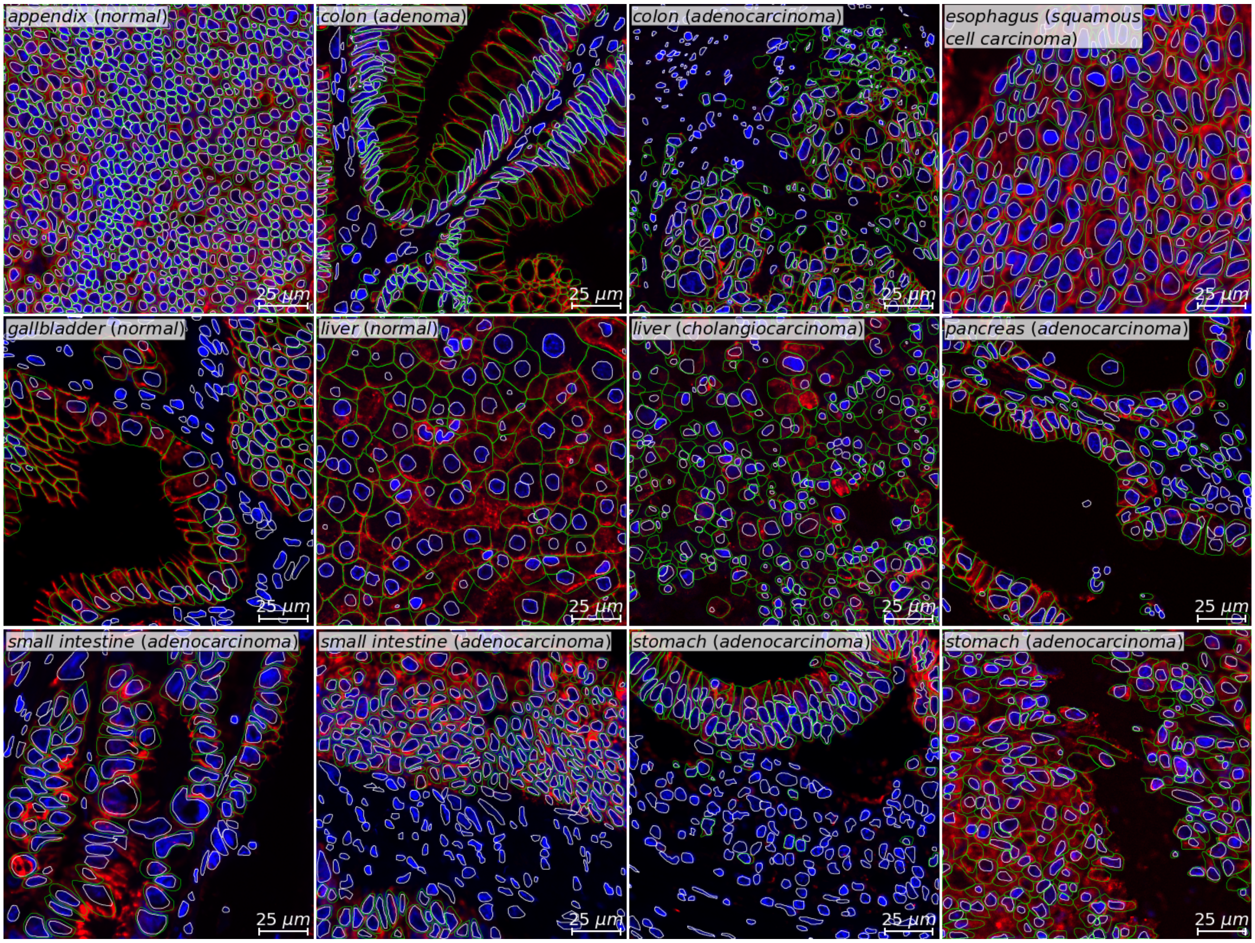


Supplementary Figure 2: Twelve images from the GIT dataset corresponding to the images in Figure 3 of the main manuscript, with expert-annotated nuclei (outlined in white) and cells (outlined in green). Blue and red colors respectively indicate nucleus (Hoechst) and cell membrane ($Na^{+}K^{+}$ATPase) marker expressions.

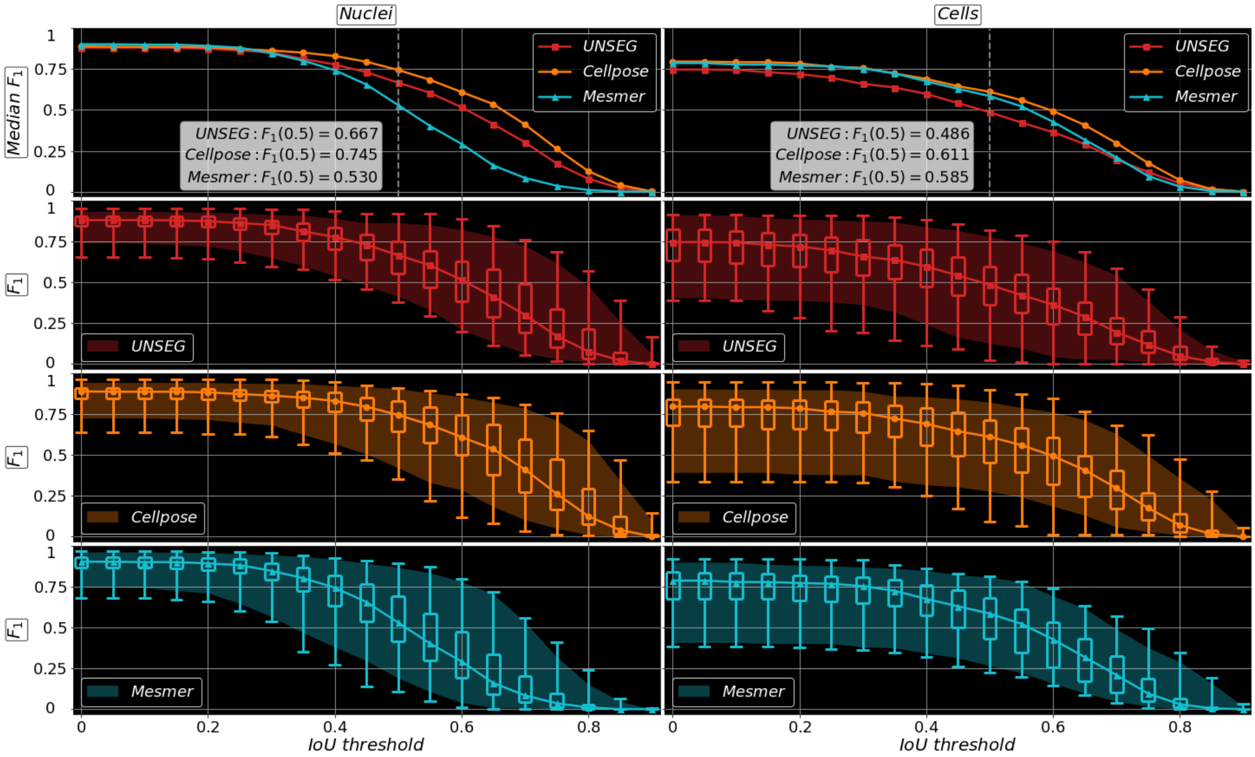


Supplementary Figure 3: Detailed performance comparison of UNSEG, Cellpose, and Mesmer on the GIT dataset. This figure supplements Figure 5 in the main manuscript. The first row is the same as the first row of Figure 5. The second, third, and fourth rows show the median $F_1$ score curves, 95% confidence intervals, and box-and-whisker plots computed for UNSEG, Cellpose, and Mesmer, respectively, as functions of IoU threshold for nuclei (left panel) and cells (right) panel. The box and whisker plots display the $F_1$ score range and the interquartile range.

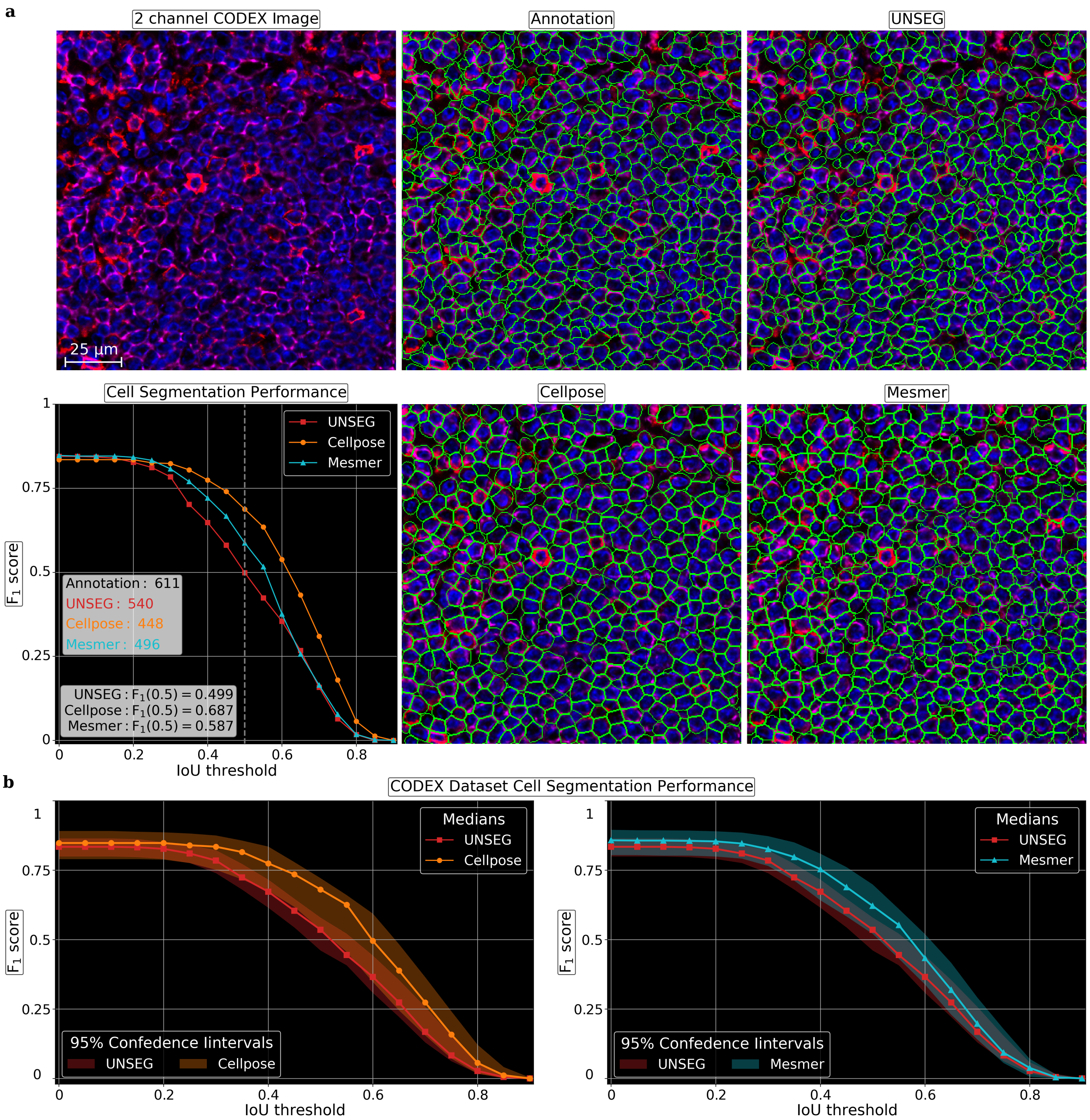


Supplementary Figure 4: Cell segmentation performance comparison of UNSEG, Cellpose, and Mesmer on annotated dataset acquired using CODEX imaging platform. **a** Six panels displayed clockwise from the top left panel to bottom left. First panel shows one representative 2-channel image of a lymph node with densely packed cells, where blue (DAPI) and red (CD20 and CD45RO) colors indicate nucleus and cell membrane markers, respectively. Second panel shows the image with annotated cells (shown in green). Third, fourth, and fifth panels respectively show the image with cells (shown in green) segmented by UNSEG, Mesmer, and Cellpose. Sixth panel shows cell segmentation accuracy of UNSEG, Cellpose, and Mesmer, measured using number of segmented cells, and $F_1$ score curves plotted as a function of IoU threshold. **b** The left and right panels respectively show pairwise cell segmentation comparison between UNSEG and Cellpose, and UNSEG and Mesmer for the entire CODEX dataset. They show median $F_1$ score curves along with their 95% confidence intervals as a function of IoU threshold. Only the following UNSEG parameters were modified from their default values indicated in Supplementary Table 2: $d_0 = 1.7$, $u_0 = 2$, $r_0 = 3$, and $n_0 = \{2, 5, 10, 20, 40\}$.

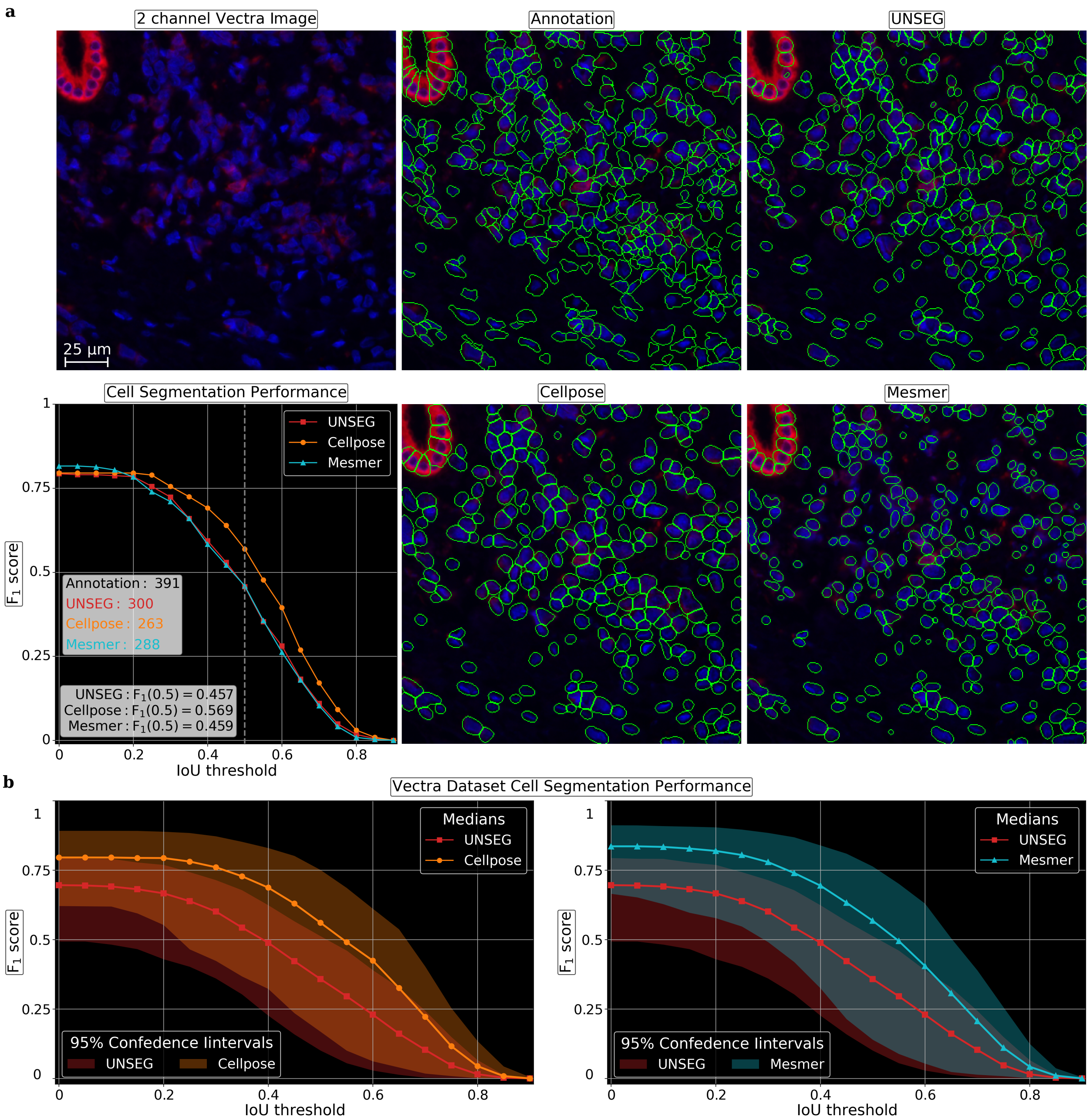


Supplementary Figure 5: Cell segmentation performance comparison of UNSEG, Cellpose, and Mesmer on annotated dataset acquired using Vectra imaging platform. **a** Six panels displayed clockwise from the top left panel to bottom left. First panel shows one representative 2-channel image of a pancreatic ductal adenocarcinoma sample, where blue (DAPI) and red (pan-Cytokeratin) colors indicate nucleus and cell membrane markers, respectively. Second panel shows the image with annotated cells (shown in green). Third, fourth, and fifth panels respectively show the image with cells (shown in green) segmented by UNSEG, Mesmer, and Cellpose. Sixth panel shows cell segmentation accuracy of UNSEG, Cellpose, and Mesmer, measured using number of segmented cells, and $F_1$ score curves plotted as a function of IoU threshold. **b** The left and right panels respectively show pairwise cell segmentation comparison between UNSEG and Cellpose, and UNSEG and Mesmer for the entire Vectra dataset. They show median $F_1$ score curves along with their 95% confidence intervals as a function of IoU threshold. Only the following UNSEG parameters were modified from their default values indicated in Supplementary Table 2: $d_0 = 1.5$, $u_0 = 2$, $r_0 = 3$, and $n_0 = \{2, 5, 10, 20, 40\}$.

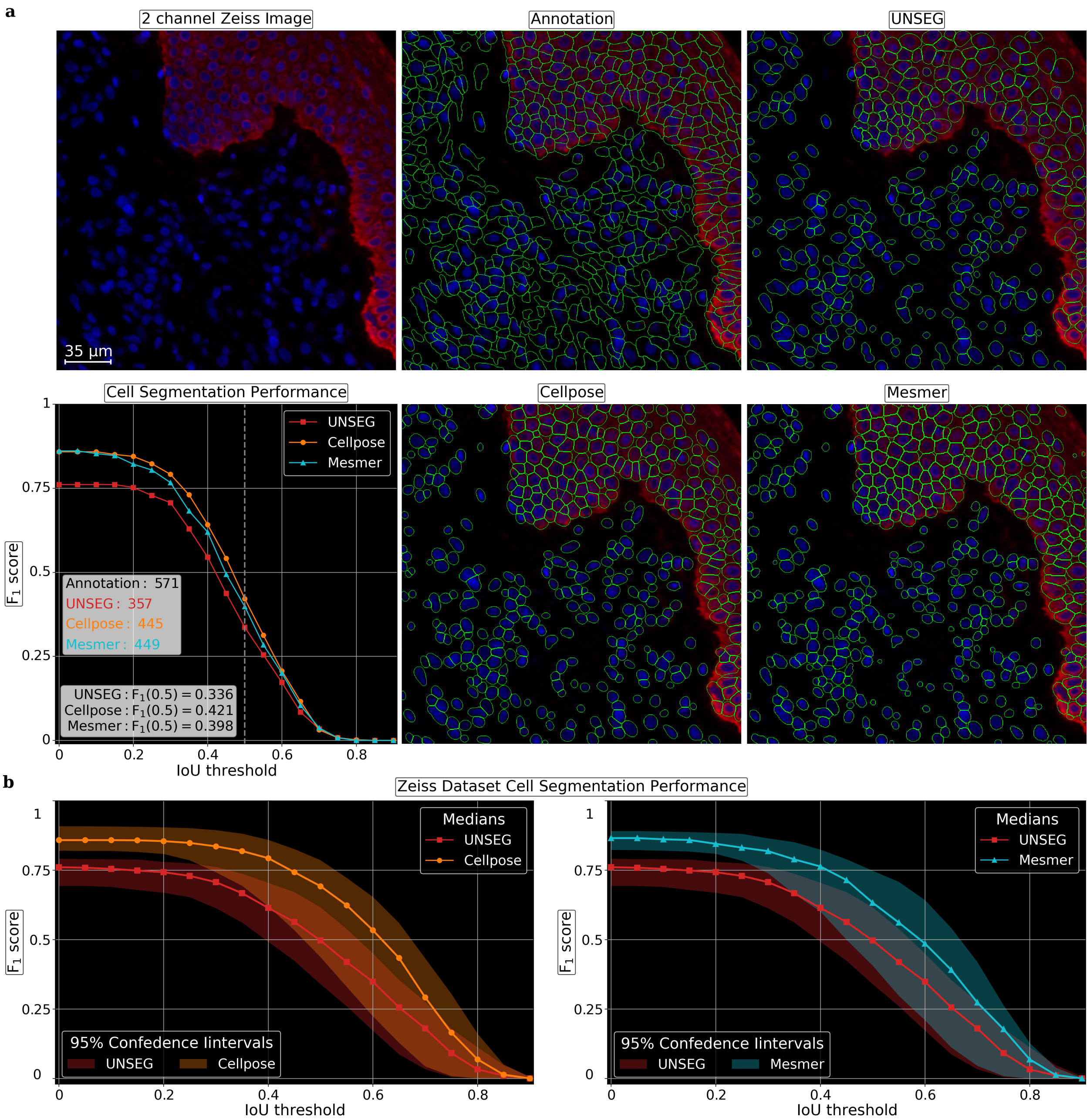


Supplementary Figure 6: Cell segmentation performance comparison of UNSEG, Cellpose, and Mesmer on annotated dataset acquired using Zeiss imaging platform. **a** Six panels listed clockwise from the top left panel to bottom left. First panel shows one representative 2-channel image, where blue and red colors indicate nucleus and cell membrane markers, respectively. Second panel shows the image with annotated cells (shown in green). Third, fourth, and fifth panels respectively show the image with cells (shown in green) segmented by UNSEG, Mesmer, and Cellpose. Sixth panel shows cell segmentation accuracy of UNSEG, Cellpose, and Mesmer, measured using number of segmented cells, and $F_1$ score curves plotted as a function of IoU threshold. **b** The left and right panels respectively show pairwise cell segmentation comparison between UNSEG and Cellpose, and UNSEG and Mesmer for the entire Zeiss dataset. They show median $F_1$ score curves along with their 95% confidence intervals as a function of IoU threshold. Only the following UNSEG parameters were modified from their default values indicated in Supplementary Table 2: $d_0 = 1.7$, $u_0 = 2$, $r_0 = 3$, and $n_0 = \{2, 5, 10, 20, 40\}$.

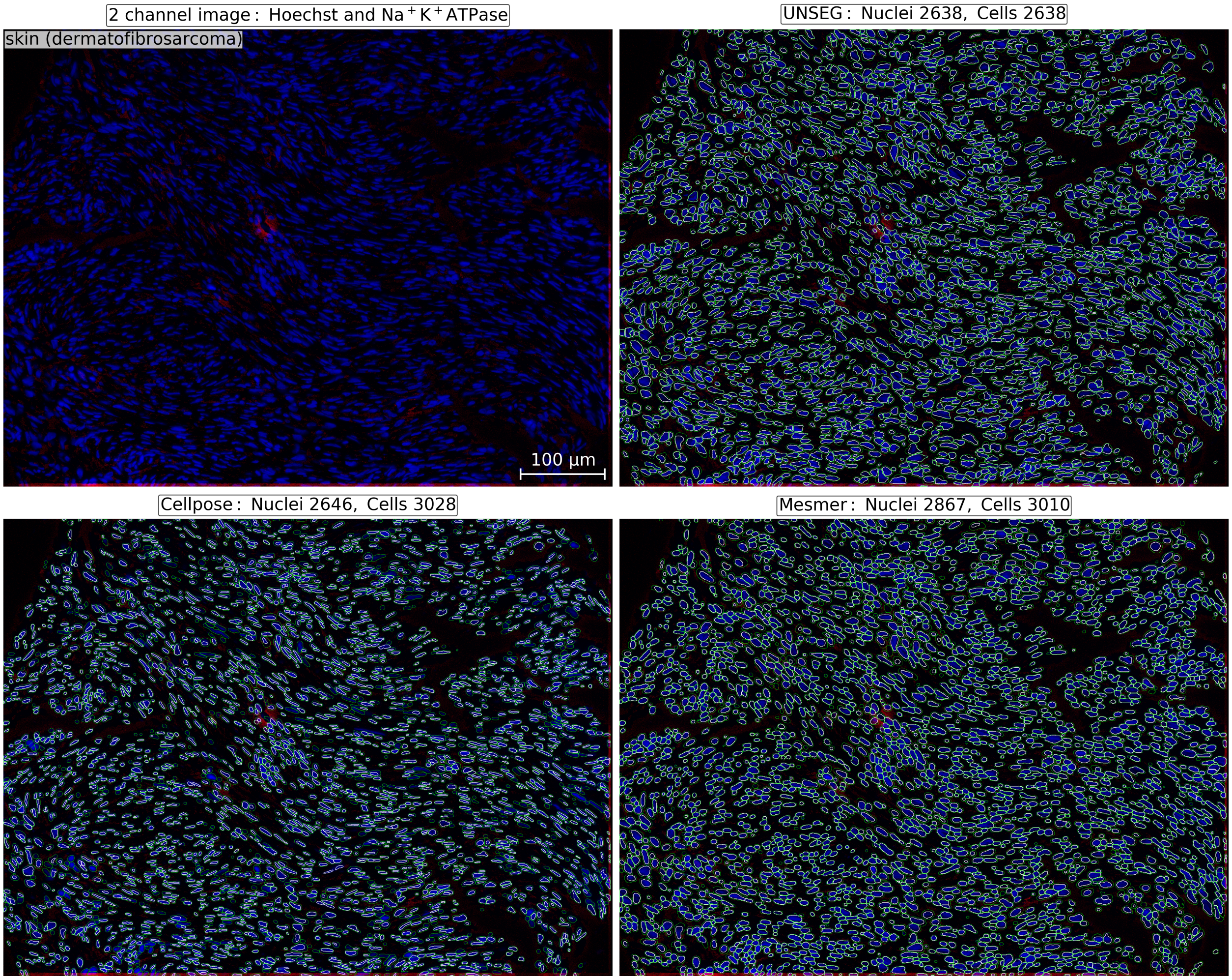


Supplementary Figure 7: Visual comparison of UNSEG, Cellpose, and Mesmer segmentation performance on an immunofluorescence image with a weak cell membrane marker expression. Four panels displayed clockwise from the top left panel to bottom left. First panel shows 2-channel image of skin tissue with dermatofibrosarcoma, where blue (Hoechst) and red ($Na^+K^+$ATPase) colors indicate nucleus and cell membrane markers, respectively. As can be seen $Na^+K^+$ATPase is weakly expressed. Second, third, and fourth panels respectively show the image with nuclei (shown in white) and cells (shown in green) segmented by UNSEG, Mesmer, and Cellpose. These panels also show the total number of nuclei and cells segmented by the methods. Only the following UNSEG parameters were modified from their default values indicated in Supplementary Table 2: $d_0 = 1.8$, and $u_0 = 4$.

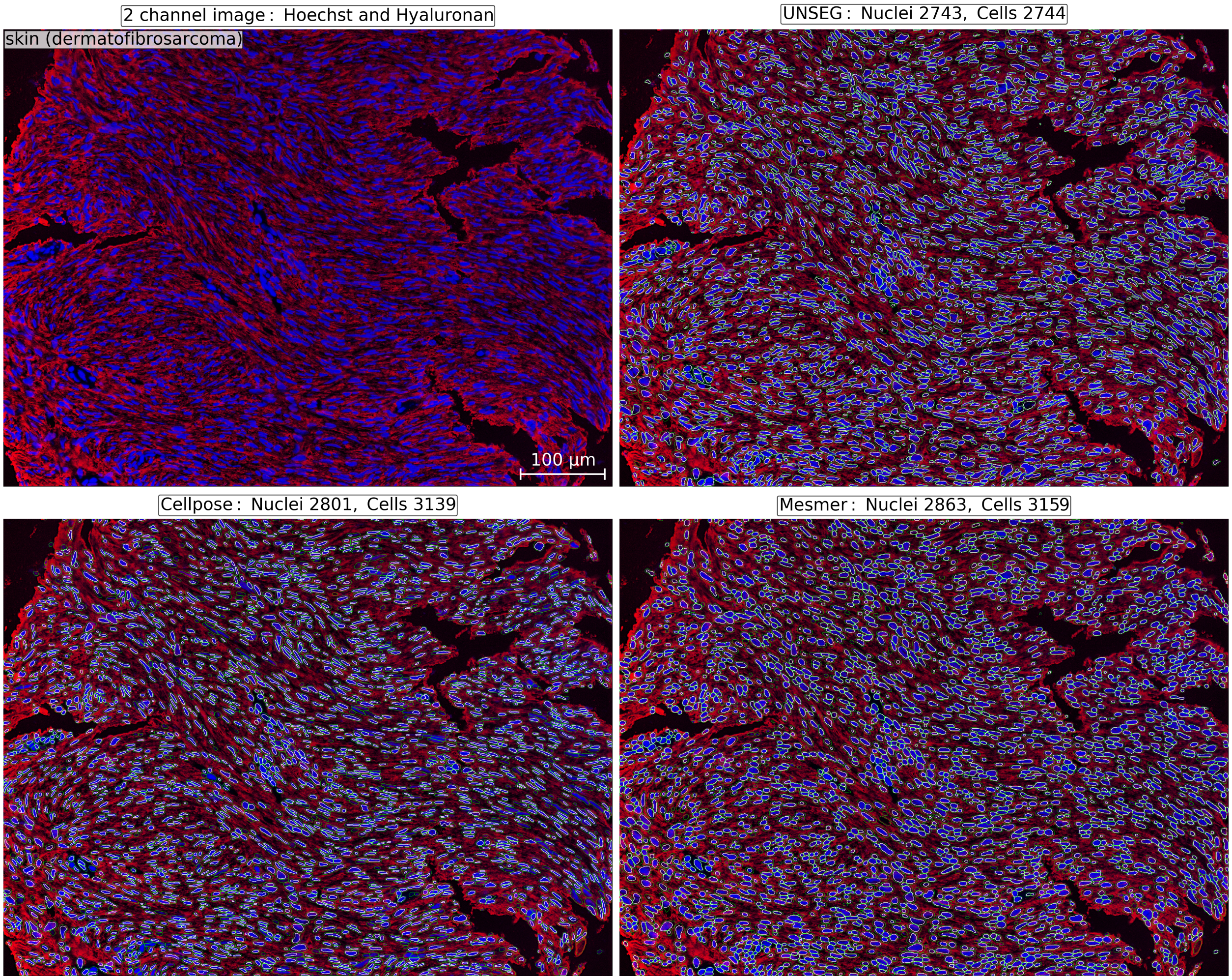


Supplementary Figure 8: Visual comparison of UNSEG, Cellpose, and Mesmer segmentation performance on an immunofluorescence image with a strong, non-specific cell-membrane marker expression. Four panels displayed clockwise from the top left panel to bottom left. First panel shows 2-channel image of skin tissue with dermatofibrosarcoma, where blue (Hoechst) and red (Hyaluronan) colors indicate nucleus and non-specific cell-membrane markers, respectively. Here, Hyaluronan not only localizes to the cell membrane but also to the cytoplasm and the extra-cellular matrix. Second, third, and fourth panels respectively show the image with nuclei (shown in white) and cells (shown in green) segmented by UNSEG, Mesmer, and Cellpose. These panels also show the total number of nuclei and cells segmented by the methods. Only the following UNSEG parameters were modified from their default values indicated in Supplementary Table 2: $d_0 = 2.5$ and $u_0 = 4$.

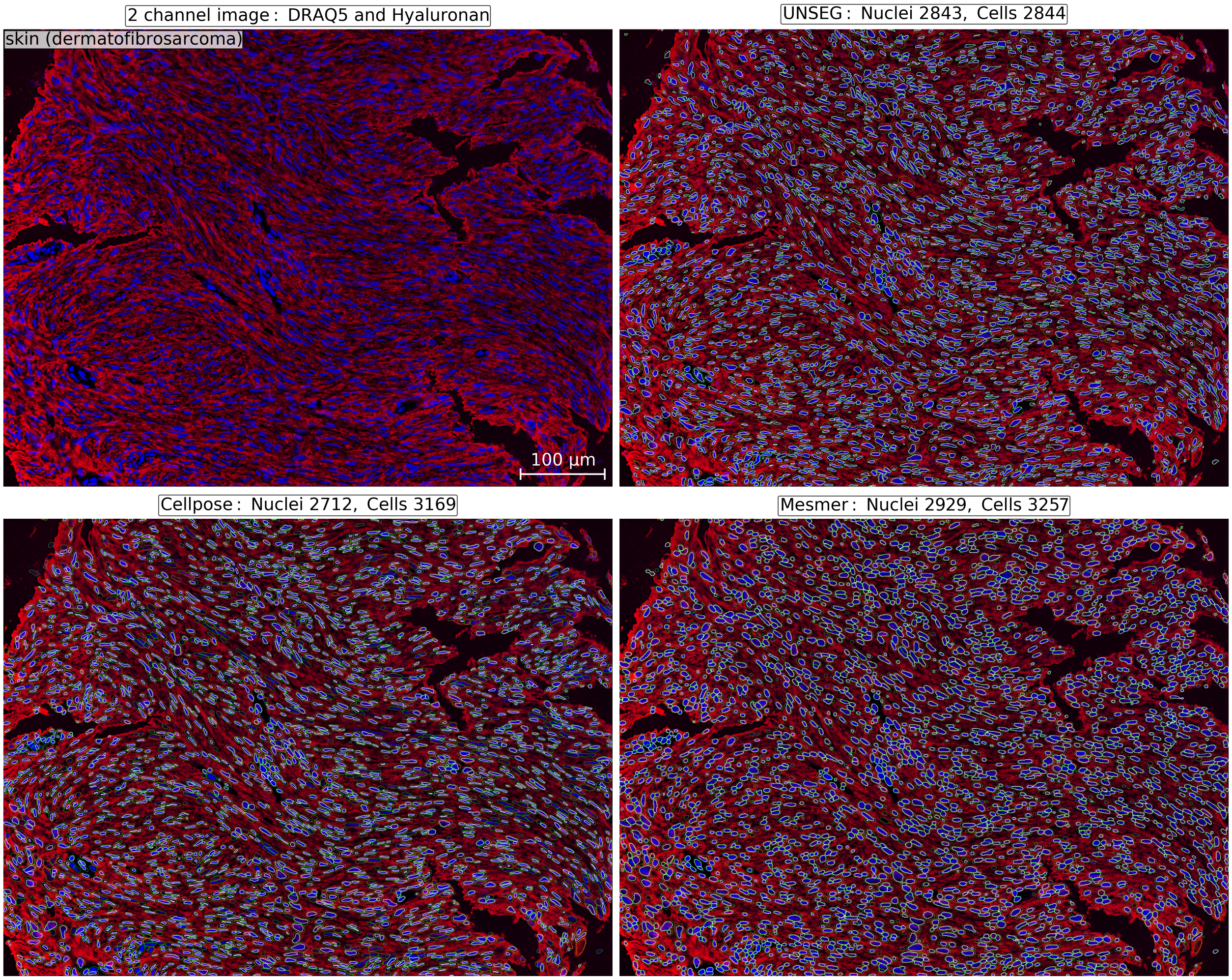


Supplementary Figure 9: Visual comparison of UNSEG, Cellpose, and Mesmer segmentation performance on an immunofluorescence image with DRAQ5 as the nucleus marker instead of Hoechst. Four panels displayed clockwise from the top left panel to bottom left. First panel shows 2-channel image of skin tissue with dermatofibrosarcoma, where blue (DRAQ5) and red (Hyaluronan) colors indicate nucleus and non-specific cell membrane markers, respectively. Second, third, and fourth panels respectively show the image with nuclei (shown in white) and cells (shown in green) segmented by UNSEG, Mesmer, and Cellpose. These panels also show the total number of nuclei and cells segmented by the methods. Only the following UNSEG parameters were modified from their default values indicated in Supplementary Table 2: $d_0 = 2.5$ and $u_0 = 4$.

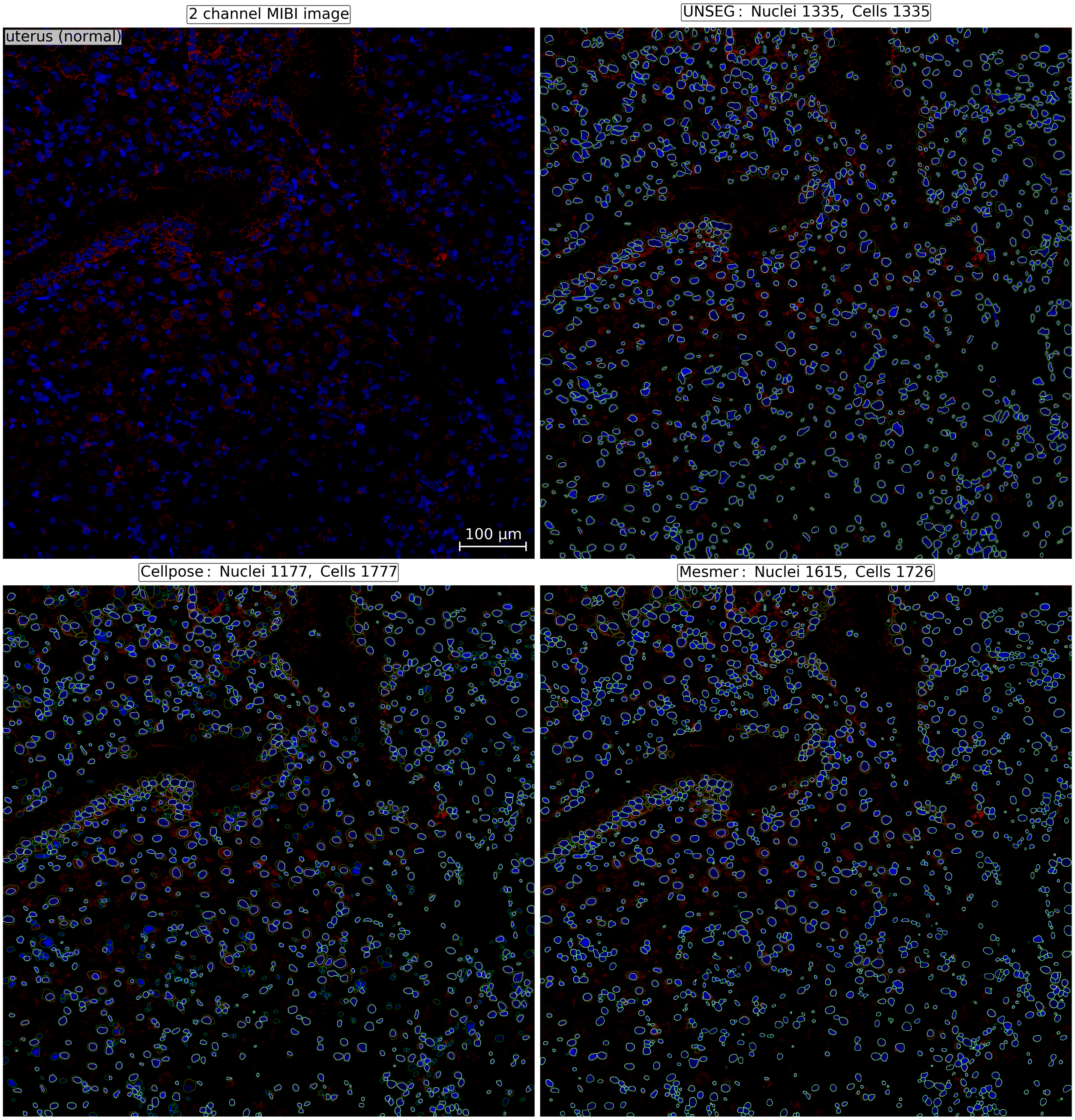


Supplementary Figure 10: Visual comparison of UNSEG, Cellpose, and Mesmer segmentation performance on a MIBI image with nucleus and cell membrane markers. Four panels displayed clockwise from the top left panel to bottom left. First panel shows 2-channel MIBI image of placental tissue, where blue and red colors indicate nucleus and cell membrane markers, respectively. Second, third, and fourth panels respectively show the image with nuclei (shown in white) and cells (shown in green) segmented by UNSEG, Mesmer, and Cellpose. Second, third, and fourth panels also show the total number of nuclei and cells segmented by the methods. Only the following UNSEG parameters were modified from their default values indicated in Supplementary Table 2: $a_0 = 5$, $d_0 = 3.5$, and $u_0 = 4$.

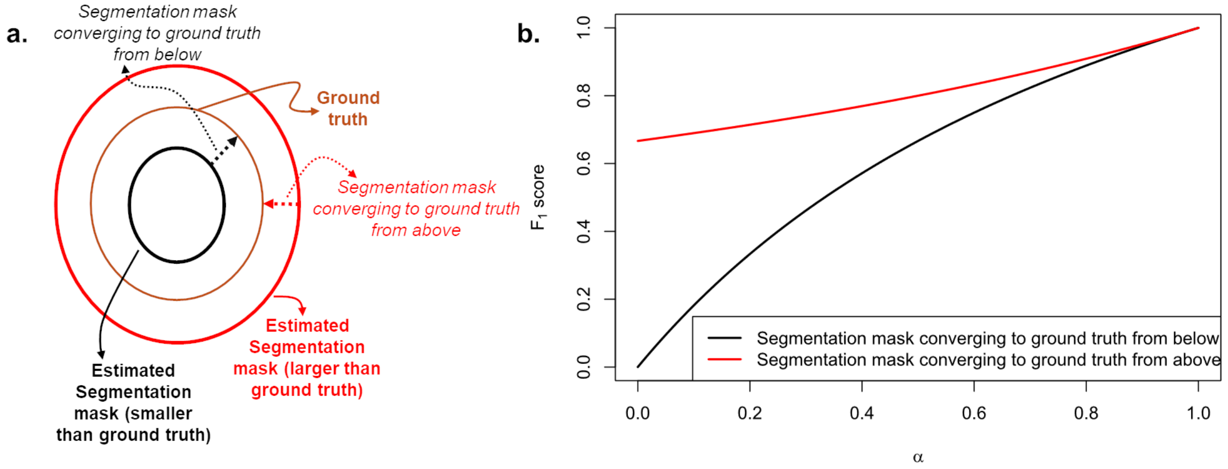


Supplementary Figure 11: Comparing quantitative accuracy of $F_1$ score in over and undersegmented scenarios. **a** Stylized depiction of a ground truth mask in brown, with under- (red) and over- (black) segmented mask estimates. Here, under-segmentation implies the segmentation mask size is larger than the ground truth, while over-segmentation implies segmentation mask is smaller than the mask size. Given these masks, **b**, illustrates how $F_1$ score behaves as the under- (red curve) and over- (black curve) segmented masks converge to the ground truth respectively from above and below. This convergence is visually shown respectively by dashed red and black arrows in (**a**). Mathematically, the $F_1$ score for convergence from above is given by $\frac{2}{3-\alpha}, \alpha \in [0, 1]$. When convergence is from below, $F_1$ is given by $\frac{2\alpha}{1+\alpha}, \alpha \in [0, 1]$. The two curves demonstrate that convergence from below is upper bounded by convergence from above. As a result, is shows that if the size of the estimated mask is larger than the ground truth, then the $F_1$ score would be higher than when the estimated mask is smaller. Therefore, even when both estimates, strictly speaking, are incorrect, one gives a better $F_1$ score than the other.

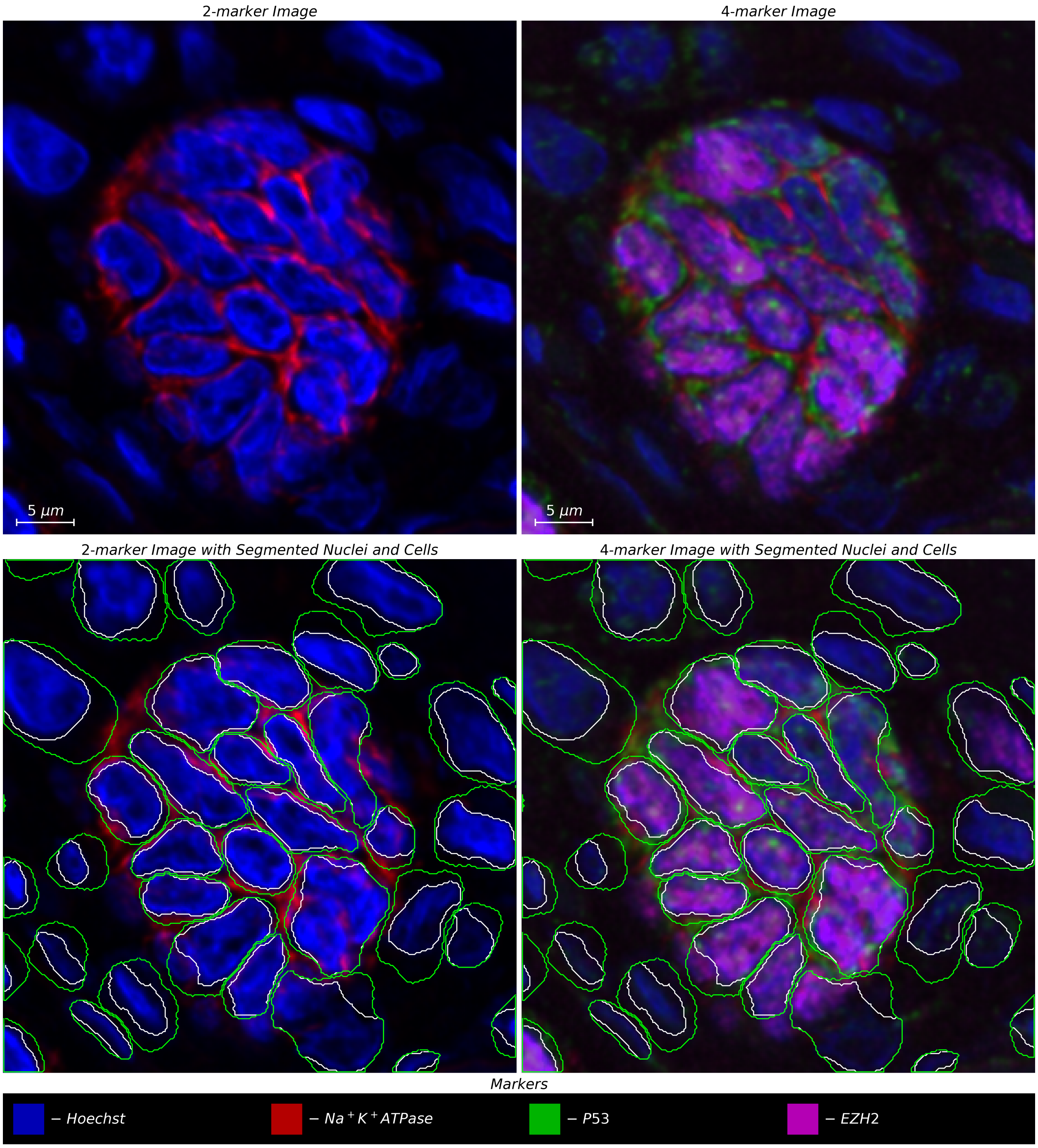


Supplementary Figure 12: UNSEG use case. Multiplexed image of healthy colon tissue from a region with densely packed cells labeled with Hoechst, $Na^{+}K^{+}$ATPase, P53, and EZH2 markers (top right). Hoechst and $Na^{+}K^{+}$ATPase channels (top left) are extracted by UNSEG to perform nucleus and cell segmentation (bottom left). When applied to the multiplexed image, the internally consistent UNSEG segmentation boundaries of cells (shown in green) and their nuclei (shown in white) are able to correctly localize sub-cellular P53, and EZH2 expression. Thus, UNSEG can be used to study intra-cellular signaling, where cellular localization of signaling pathway components is required to be correctly identified.

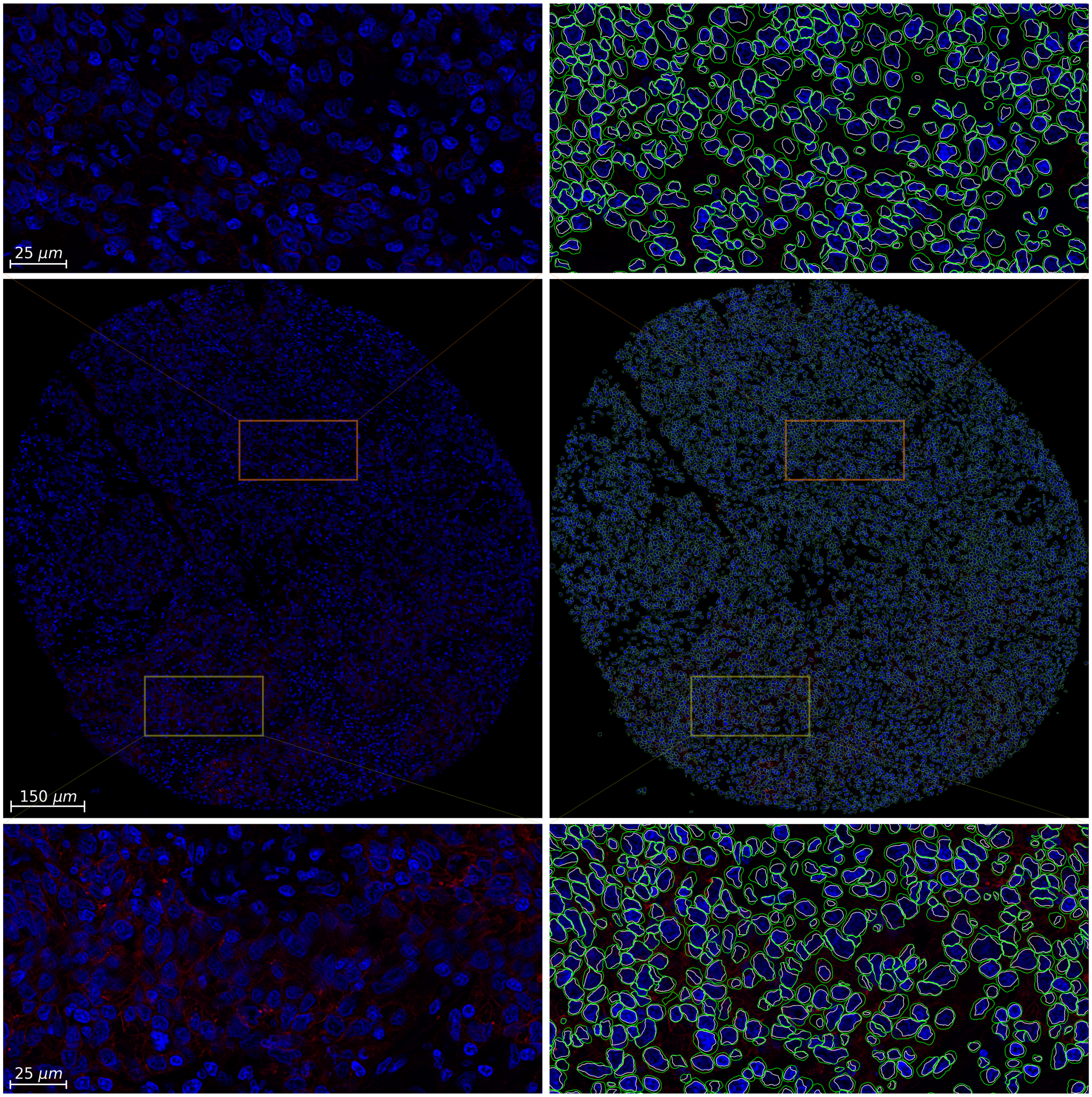


Supplementary Figure 13: UNSEG based nucleus and cell segmentation of a 1mm colon tissue microarray spot. The image size is 6850 × 6850 pixels, the pixel pitch of 0.16$\mu$m/pixel. This image was also used to compute the runtime complexity of UNSEG.

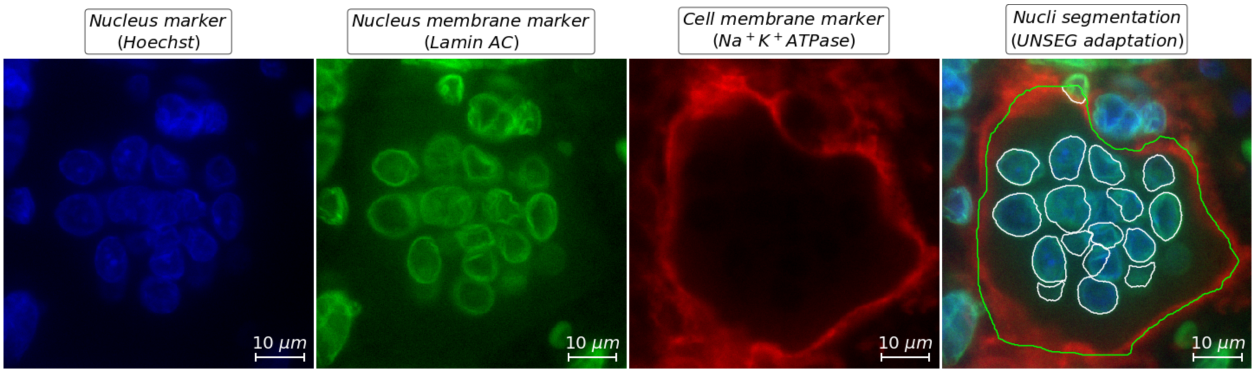


Supplementary Figure 14: Segmentation of a multi-nucleated cell from a tissue section of giant cell tumor of bone, obtained using UNSEG modification based joint unsupervised processing of nucleus marker (Hoechst) and nucleus membrane marker (Lamin A/C), with cell membrane marker used for identifying the cell membrane. The contours of segmented nuclei and cell are shown in white and green, respectively.

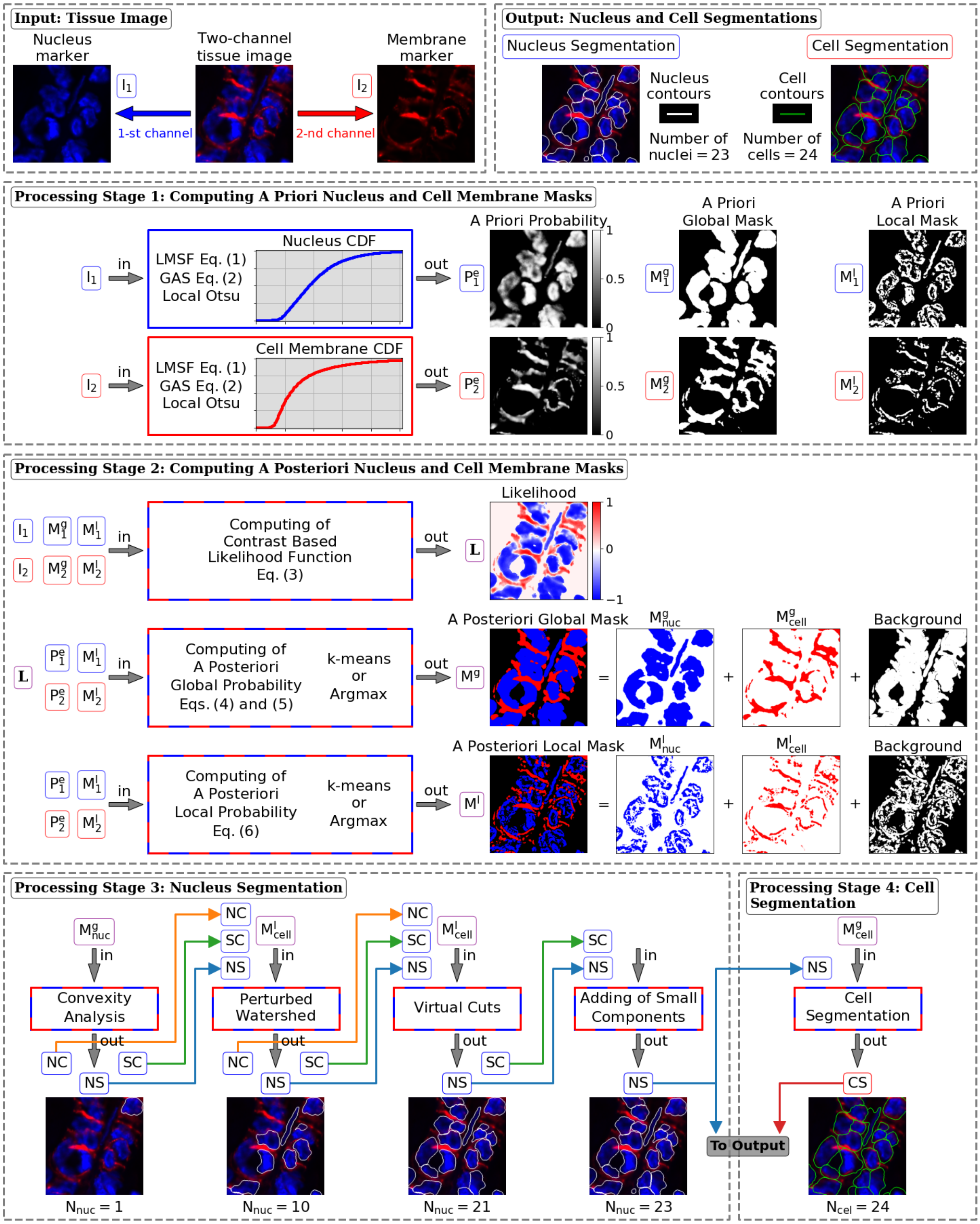


Supplementary Figure 15: Systematic illustration of the four processing stages of UNSEG detailed in the Methods section. NC and SC indicate two lists of nucleus clusters and small components. NS and CS stand for nucleus and cell segmentations, respectively.

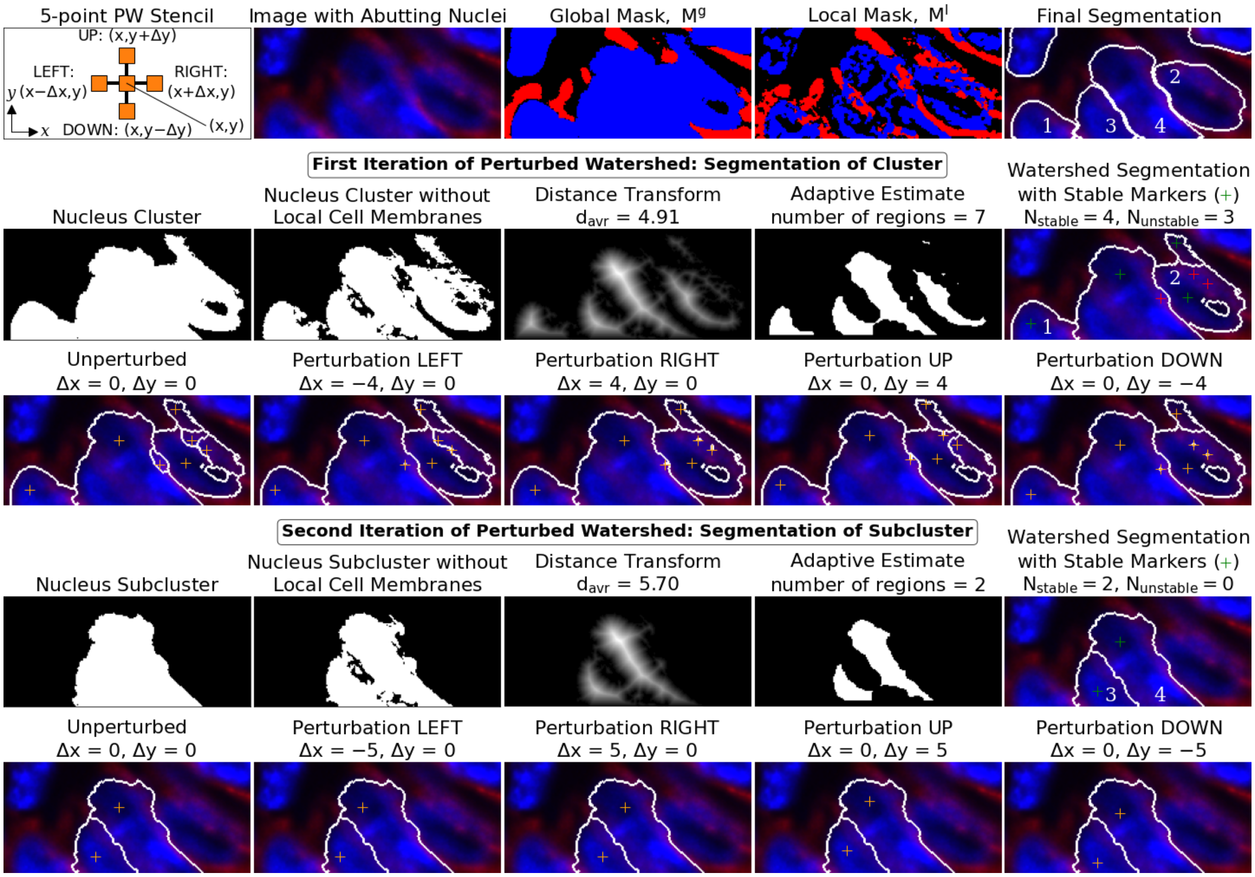


Supplementary Figure 16: Detailed illustration of nested perturbed watershed. The first iteration shows that the original nucleus cluster was split into four objects associated with four stable seed points (the last panel in the second row). Two of them (labeled 1 and 2) were classified as nuclei, the other two were classified respectively as a sub-cluster (the largest object) and a small object (the smallest object). The second iteration shows that the sub-cluster was split into two objects associated with two stable seed points (the last panel in the fourth row). Both objects (labeled 3 and 4) were classified as nuclei. Thus, the final perturbed watershed segmentation consists of four nuclei (the last panel in the first row). The panels in the first row (from left to right) present the perturbed watershed stencil (defines the four directions of displacements of watershed markers in perturbed iterations, where $\Delta x = \Delta y = \lfloor d_{avr} \rfloor$ pixels), the fragment of the two-channel image with nucleus cluster, *a posteriori* global and local masks (input), and the final nucleus segmentation of cluster based on the nested perturbed watershed (output). The second and third rows illustrate the first iteration of the perturbed watershed, where the four gray-scale images demonstrate the preparatory calculations. The fourth and fifth rows do the same for the second iteration. The color images in the third and fifth rows show the unperturbed and four perturbed segmentation possibilities respectively for each iteration.
